# Early acquisition of conserved, lineage-specific proteins currently lacking functional predictions were central to the rise and diversification of archaea

**DOI:** 10.1101/2020.07.16.207365

**Authors:** Raphaël Méheust, Cindy J. Castelle, Alexander L. Jaffe, Jillian F. Banfield

**Affiliations:** Department of Earth and Planetary Science, University of California, Berkeley, CA; Chan Zuckerberg Biohub, San Francisco, CA; Department of Environmental Science, Policy, and Management, University of California, Berkeley, CA; Innovative Genomics Institute, University of California, Berkeley, CA; Department of Plant and Microbial Biology, University of California, Berkeley, CA

## Abstract

Recent genomic analyses of Archaea have profoundly reshaped our understanding of their distribution, functionalities and roles in eukaryotic evolution. Within the domain, major supergroups are Euryarchaeota, which includes many methanogens, the TACK, which includes Thaumarchaeaota that impact ammonia oxidation in soils and the ocean, the Asgard, which includes lineages inferred to be ancestral to eukaryotes, and the DPANN, a group of mostly symbiotic small-celled archaea. Here, we investigated the extent to which clustering based on protein family content recapitulates archaeal phylogeny and identified the proteins that distinguish the major subdivisions. We also defined 10,866 archaeal protein families that will serve as a community resource. Clustering based on these families broadly recovers the archaeal phylogenetic tree. Interestingly, all major groups are distinguished primarily by the presence of families of conserved hypothetical proteins that are either novel or so highly diverged that their functions are obscured. Given that these hypothetical proteins are near ubiquitous within phyla, we conclude that they were important in the origin of most of the major archaeal lineages.

## Introduction

Until recently, the archaeal domain comprised only two phyla, the Euryarchaeota and the Crenarchaeota, most of which were described from extreme environments (Woese, Kandler, and Wheelis 1990; Woese and Fox 1977). The recovery of genomes from metagenomes without the prerequisite of laboratory cultivation has altered our view of diversity and function across the Archaea domain (Spang, Caceres, and Ettema 2017; Adam et al. 2017; Baker et al. 2020). Hundreds of genomes from little studied and newly discovered archaeal clades have provided new insights into archaeal metabolism and evolution. Now, Archaea include at least four major large groups, the Euryarchaeota (Cluster I and Cluster II; (Spang, Caceres, and Ettema 2017; Adam et al. 2017; Baker et al. 2020)), the TACK (Proteoarchaeota) (Petitjean et al. 2014), the Asgard (Spang et al. 2015; Zaremba-Niedzwiedzka et al. 2017), and the DPANN (Castelle et al. 2015; Rinke et al. 2013), all of which comprise several distinct phylum-level lineages. These archaea are not restricted to extreme habitats, but are widely distributed in diverse ecosystems (Spang, Caceres, and Ettema 2017; Adam et al. 2017; Baker et al. 2020).

Most studies have focused on the metabolic potential of archaea based on analysis of proteins with known functions and revealed roles in the carbon, nitrogen, hydrogen and sulfur biogeochemical cycles. For example, Euryarchaeota includes many methanogens and non-methanogens, including heterotrophs and sulfur oxidizers (Offre, Spang, and Schleper 2013). The TACK includes Thaumarchaeota, most but not all of which oxidize ammonia (Pester, Schleper, and Wagner 2011; Brochier-Armanet et al. 2008), Aigarchaeota that tend to be chemolithotrophs that oxidize reduced sulfur compounds (Hua et al. 2018), Crenarchaeota that include thermophilic sulfur oxidizers (Woese et al. 1984), and Korarchaeota, a highly undersampled group represented by amino acid degraders that anaerobically oxidize methane and also metabolize sulfur compounds (McKay et al. 2019). The Asgard have variable metabolisms and their genomes encode pathways involved in structural components that are normally considered to be eukaryotic signatures (Spang et al. 2015; Zaremba-Niedzwiedzka et al. 2017). The DPANN are an intriguing group that typically have very small genomes and symbiotic lifestyles (Castelle et al. 2018; Dombrowski et al. 2019). Their geochemical roles are difficult to predict, given the predominance of hypothetical proteins.

Previously, the distribution of protein families over bacterial genomes was used to provide a function rather than phylogeny-based clustering of lineages (Méheust et al. 2019). Protein clustering allows the comparison of the gene content between genomes by converting amino acid sequences into units of a common language. The method is agnostic and unbiased by preconceptions about the importance or functions of genes.

Here, we adapted this approach to evaluate the protein family-based coherence of the archaea and to test the extent to which a subdivision of archaea could be resolved based on shared protein family content. The analysis drew upon the large genome dataset that is now available for cultivated as well as uncultivated archaea (3,197 genomes). The observation that hypothetical proteins dominate the sets of co-occurring protein families that distinguish major archaeal groups indicates the importance of these protein sets in the rise of the major archaeal lineages.

## Results

### Genome reconstruction and collection improved the resolution of the DPANN lineages

We collected 2,618 genomes spanning all the recognized phyla and superphyla of the Archaea domain from the NCBI genome database (**Supplemental Dataset – Table S1**). To enable our analyses, we augmented the relatively limited sampling of the DPANN by adding 569 newly available DPANN draft genomes (Castelle et al. in prep.) from low oxygen marine ecosystems, an aquifer adjacent to the Colorado River, Rifle, Colorado, and from groundwater collected at the Genasci dairy farm, Modesto CA (He et al., n.d.). The 3,197 genomes were clustered at ≥ 95% average nucleotide identity (ANI) to generate 1749 clusters. We removed genomes with <70% completeness or >10% contamination or if there was < 50% of the expected columns in the alignment of 14 concatenated ribosomal proteins (**see Materials and Methods**). To avoid contamination due to mis-binning, we required that these proteins were co-encoded on a single scaffold. The average completeness of the final set of 1,179 representative genomes is 95% and 928 were >90% complete (**Supplementary Dataset – Table S1).** The 1,179 representative genomes comprise 39 phylum-level lineages including 16 phyla that have more than 10 genomes (**Supplementary Dataset – Table S1 and Figure 1**).

**Figure 1.**
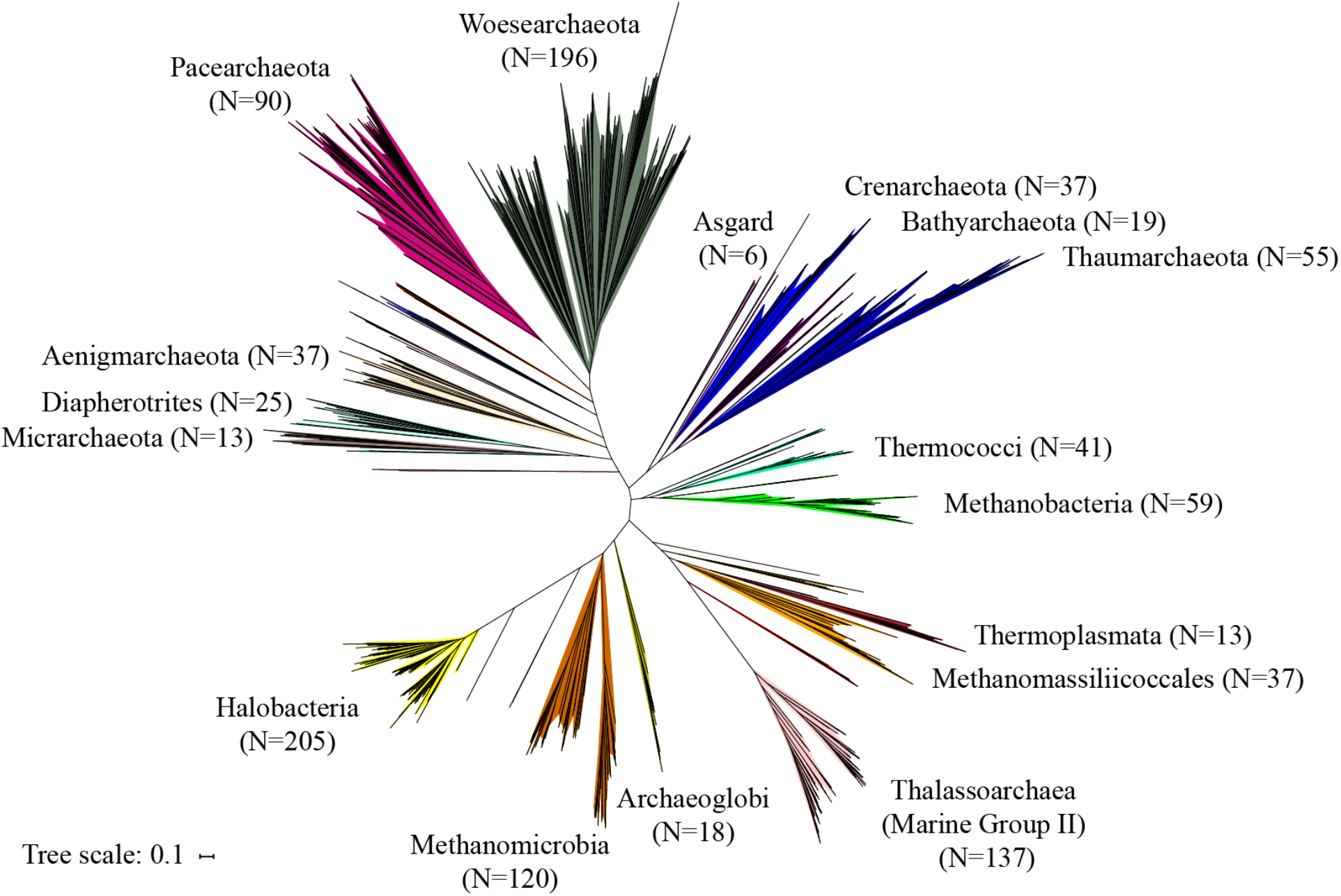
Phylogenetic tree of the 1,179 representative genomes. The maximum-likelihood tree was calculated based on the concatenation of 14 ribosomal proteins (L2, L3, L4, L5, L6, L14, L15, L18, L22, L24, S3, S8, S17, and S19) using the LG plus gamma model of evolution. Scale bar indicates the average substitutions per site. The complete ribosomal protein tree is available in rectangular format in Supplementary Figure 1.

### Genomic content of representative genomes correlates with the phylogeny of archaea

We clustered the 2,336,157 protein sequences from the representative genomes in a two-step procedure to generate groups of homologous proteins (**Supplementary Figure 2**). This resulted in 10,866 clusters (representing 2,075,863 sequences) that were present in at least five distinct genomes. These clusters are henceforth referred to as protein families.

We assessed the quality of the protein clustering. The rationale was that we expected protein sequences with the same function to cluster into the same protein family. We annotated our protein dataset using the KEGG annotations (Kanehisa et al. 2016) and systematically verified that the protein family groupings approximate functional annotations. The KEGG annotations in our dataset encompass 6,482 unique annotations of various biological processes, including fast-evolving defense mechanisms. For each of these 6,482 annotations, we reported the family that contains the highest percentage of protein members annotated with that KEGG annotation. Most clusters were of good quality. For 87% of the KEGG annotations (5,627 out of 6,482), one family always contained >80% of the proteins (**Supplementary Figure 3A**). The contamination of each protein family was assessed by computing the percentage of the proteins with KEGG annotations that differ from the dominant annotation (percentage annotation admixture). Most of the families contain only proteins with the same annotation, and 2,654 out of 3,746 families (71%) have <20% annotation admixture (**Supplementary Figure 3B**). Although this metric is useful, we note that it is imperfect because two homologous proteins can have different KEGG annotations and thus cluster into the same protein family, increasing the apparent percentage of annotation admixture. Although we used sensitive HMM-based sequence-comparison methods and assessed the quality of the protein clustering, we cannot completely rule out the possibility that our pipeline failed to retrieve distant homology for highly divergent proteins. Small proteins and fast-evolving proteins are more likely to be affected. This lack of sensitivity would result in the separation of homologous proteins into distinct families and would impact the results. To reduce the incidence of proteins without functional predictions for which annotations should have been achieved we augmented PFAM and KEGG-based annotations by comparing sequences to PDB database and by performing HMM-HMM comparison against the eggNog database (**see Materials and Methods**).

We visualized the distribution of the families over the genomes by constructing an array of the 1,179 representative genomes (rows) vs. 10,866 protein families (columns) and hierarchically clustered the genomes based on profiles of protein family presence/absence **(Figure 2A)**. The families were also hierarchically clustered based on profiles of genome presence/absence. As previously reported for bacteria (Snel, Bork, and Huynen 1999; Méheust et al. 2019), the hierarchical clustering tree of the genomes resulting from the protein clustering **(Figure 2B)** correlated with the maximum-likelihood phylogenetic tree based on the concatenation of the 14 ribosomal proteins **(Figure 1)** (the cophenetic correlation based on a complete-linkage method is 0.83, based on average-linkage 0.84, and based on single-linkage, 0.84) (**Supplementary Figure 4**). Although the tree resulting from the protein families correlates with the phylogenetic tree, it does not achieve the resolution of the phylogenetic tree, especially for placement of the deep branches. Interestingly, several phyla, such as the Crenarchaeota or the Woesarchaeota, are resolved into multiple groups (**Figure 2B**). The first clade of Woesearchaeota corresponds to the Woesarchaeota-like I whereas the second clade groups together the Woesarchaeota and Woesarchaeota-like II groups. We could not evaluate the placement of Altiarchaeota relative to the DPANN because no genomes passed our quality control thresholds.

**Figure 2.**
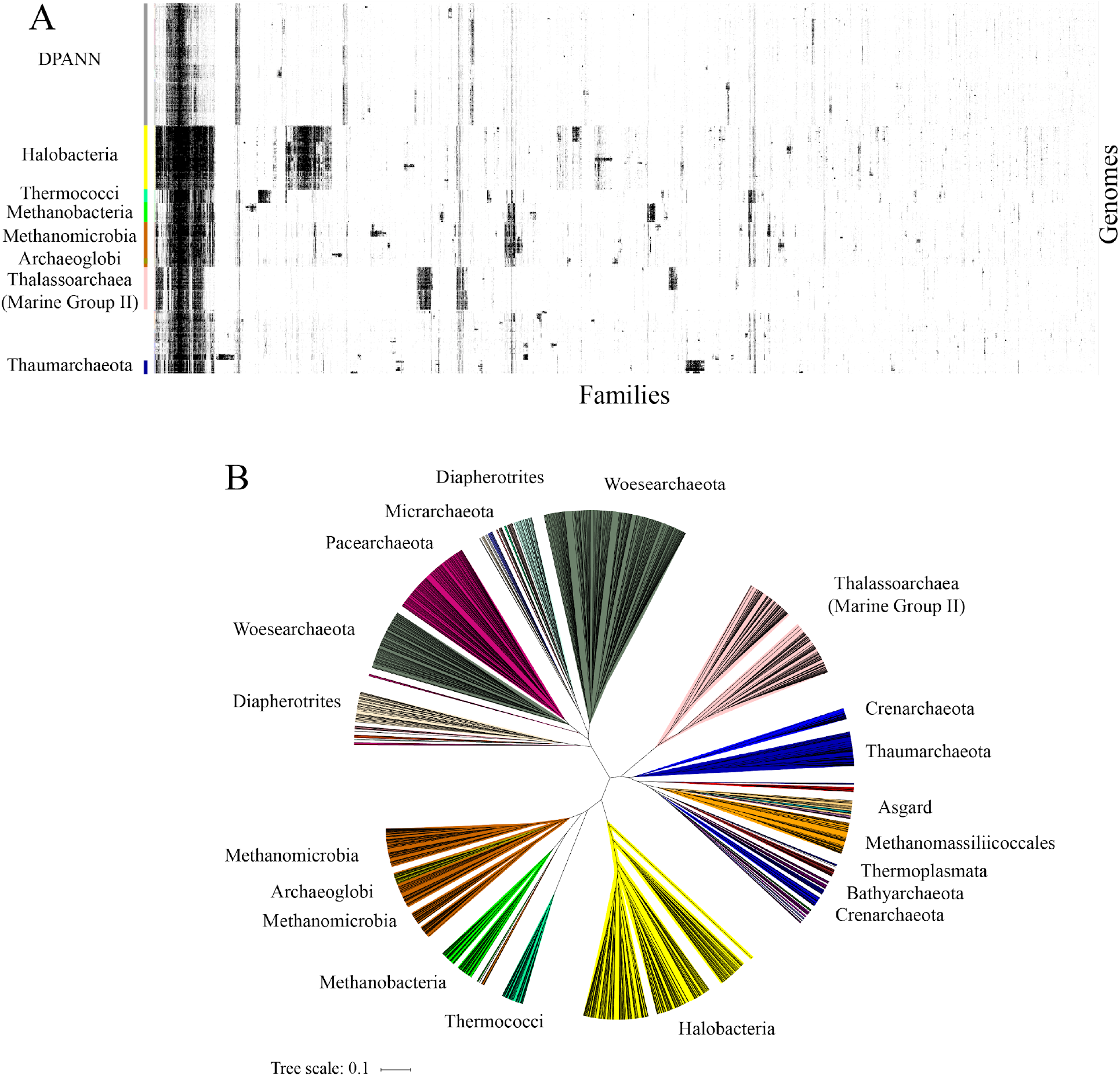
The distribution of the 10,866 families across the 1,179 representative genomes. A. The distribution of 10,866 widely distributed protein families (columns) in 1,179 representative genomes (rows) from Archaea. Data are clustered based on the presence (black) and absence (white) profiles (Jaccard distance, complete linkage). B. Tree resulting from the hierarchical clustering of the genomes based on the distributions of proteins families in panel A.

We defined modules as blocks of co-occurring protein families containing at least 20 families (see **Materials and Methods**) (Méheust et al. 2019). Each module was assigned a taxonomic distribution based on the taxonomy of the genomes with the highest number of families (see **Materials and Methods and Supplementary Dataset – Table S2**). A block of 587 protein families that was broadly conserved across the 1,179 genomes (left side in **Figure 2A**) was designed as the module of ‘core families’ (Module 1) (**Supplementary Figure 5).** Given their widespread distribution, it is unsurprising that most of the families are involved in well-known functions, including replication, transcription and translation, basic metabolism (oxidative phosphorylation chain, nucleotides, amino acids, ribosomal proteins, cofactors and vitamins, transporters, peptidases, DNA repair and chaperones). As expected, many of these easily recognized core families, primarily those involved in energy metabolism and cofactor synthesis, are absent in DPANN genomes (Castelle et al. 2018, 2015) (**Figure 2A and Supplementary Dataset – Table S3**). Another interesting module (module 23) (**Supplementary Figure 5**), composed of ~100 protein families, is widely distributed in most archaeal genomes but was not identified in DPANN and surprisingly, not in the Thalassoarchaea. Module 23 includes functions involved in carbon metabolism, amino-acid synthesis, and many transporter families. For instance, we identified several families for subunits of the Mrp antiporter as widespread in Halobacteria, Methanogens and Thermococci, but they appear to be absent in DPANN and Thalassoarchaea. The Mrp antiporter functions as Na+/H+ antiporter and also contributes to sodium tolerance in Haloarchaea. Mrp has been reported to be involved in energy conservation in methanogens and in the metabolic system of hydrogen production in Thermococci.

The DPANN are an enigmatic set of lineages, the monophyly of which remains uncertain (Aouad et al. 2018). However, the protein family analysis clearly showed that these lineages group together and are distinct from other Archaea (**Figure 2B**). A detailed protein family analysis of groups within the DPANN is presented elsewhere (Castelle et al. in prep.).

### Major non-DPANN groups possess groups of conserved protein families

We detected 96 modules that are restricted to non-DPANN lineages **(Supplementary Dataset – Table S2)**. Only 9 of the 96 modules were found in multiple phyla and in 8 of these 9 cases, the phyla that possess each module are phylogenetically unrelated (e.g., Crenarchaeota and Halobacteria). The 9th, module 44, is interesting in that it occurs in two phyla and those phyla are monophyletic (Thorarchaeota and Heimdallarchaeota of the Asgard superphylum). Thus, the vast majority of the non-DPANN modules (87) are restricted to a single phylum (**Supplementary Dataset – Table S2**) and, perhaps surprisingly given phylogenetic support for superphyla within Archaea, almost no modules are specific to superphyla.

Visualization of the distribution of protein families highlights the presence of modules that are not only lineage specific but are also well conserved within each lineage (**Figure 2A**). In fact, we identified such archaeal group-specific modules in 10 out of 11 non-DPANN with more than 10 genomes (**Table 1, Figure 3 and Supplementary Figure 5**). For instance, there are two modules (modules 13 and 108) comprising 525 families that are fairly conserved in Halobacteria. On average, each of the 525 families appears in 65% of the halobacterial genomes, yet these families are mostly absent in non-halobacterial genomes **(Supplementary Figure 6)**. These modules are slightly less conserved within each archaeal group than module 1 families (comprising core functions) (**Supplementary Figure 6)**.

**Figure 3.**
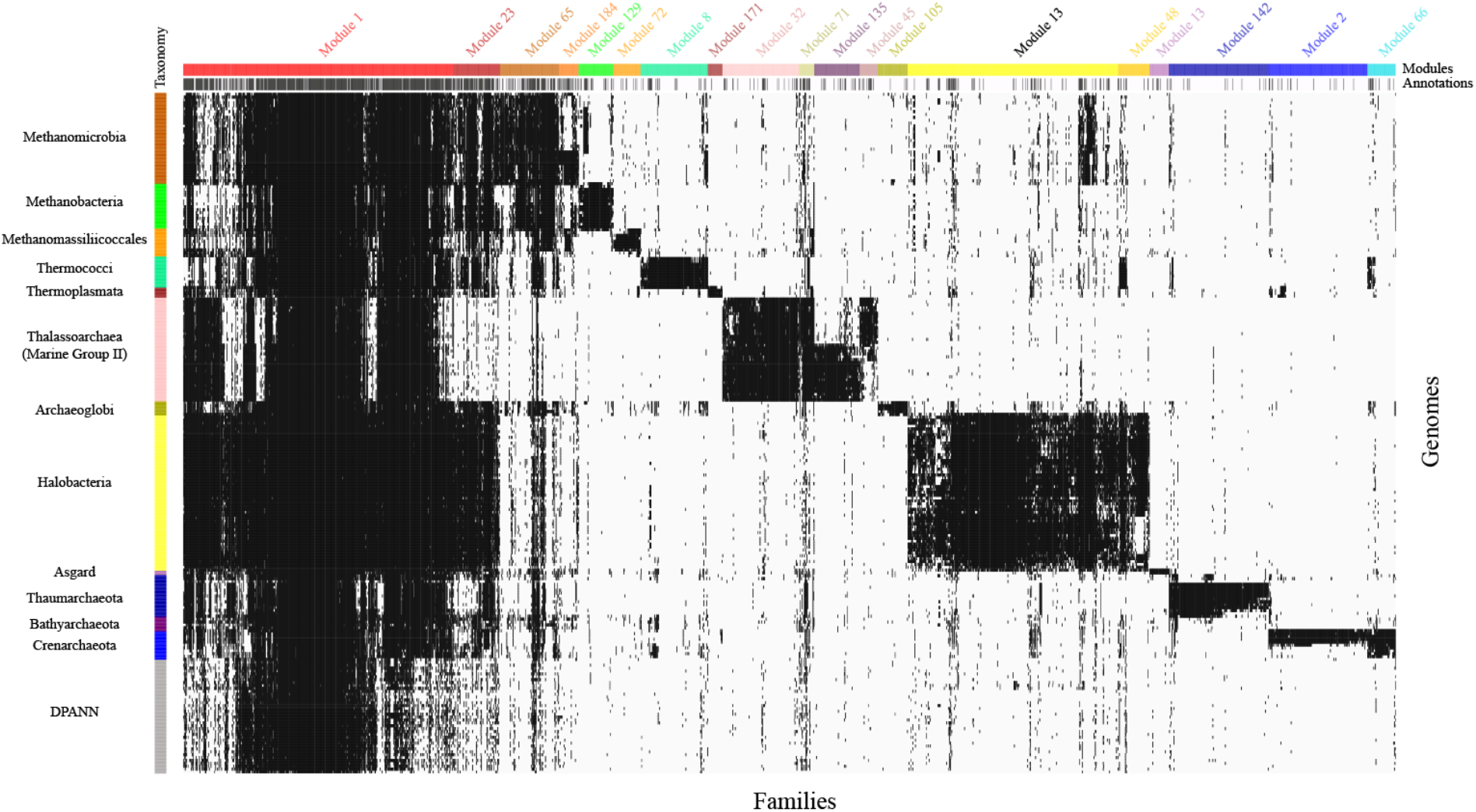
The distribution of the 2,632 families of the 19 modules discussed in this study. Each column represents a protein family and each row represents a genome. Data are clustered based on the presence (black)/ absence (white) profiles but also based on the taxonomy of the genomes and the module membership. The first colored top bar (annotations) shows the families with (black) / without (white) a predicted annotation whereas the second colored top bar (modules) indicates the module of each family. The colored side bar indicates the taxonomic assignment of each genome.

**Table 1.** **A list of the fourteen modules that are lineage specific but also well conserved within eleven major archaeal lineages. A family was counted as having a signal peptide if at least 25% of its protein sequences were predicted to have a signal peptide prediction according to the SignalP software** (Almagro Armenteros et al. 2019)**. A family was counted as having a transmembrane helix if more than half of its protein sequences were predicted to have a transmembrane helix according to the TMHMM software** (Krogh et al. 2001)**. Families were considered hypothetical if they have neither PFAM (Domain of Unknown Function domains were excluded) nor KEGG annotations (see the supplementary dataset – Table S3 for the full list of hypothetical families). Finally, a family was considered to have bacterial homologs if the family matched with protein sequences of at least ten distinct bacterial genomes (see Materials and Methods). The core module 1 is included as a comparison**.

| Modules | Lineage(s) | # Families | SignalP (%) | TMHMM (%) | Hypothetical families (%) | Hits to Bacteria (%) |
| --- | --- | --- | --- | --- | --- | --- |
| 1 | Core genome | 587 | 6 | 20 | 13 | 87 |
| 13,108 | Halobacteria | 525 | 14 | 36 | 82 | 34 |
| 66,2 | Crenarchaeota | 276 | 9 | 34 | 89 | 11 |
| 142 | Thaumarchaeota | 216 | 13 | 31 | 94 | 11 |
| 32,71 | Marine Group II | 199 | 19 | 55 | 77 | 32 |
| 8 | Thermococci | 146 | 12 | 32 | 84 | 24 |
| 65 | Methanomicrobia | 128 | 11 | 22 | 45 | 63 |
| 129 | Methanobacteria | 75 | 16 | 55 | 71 | 17 |
| 105 | Archaeoglobi | 65 | 3 | 40 | 94 | 12 |
| 72 | Methanomassiliicoccales | 59 | 22 | 49 | 86 | 27 |
| 48 | Asgard | 42 | 17 | 36 | 79 | 17 |
| 171 | Thermoplasmata | 32 | 3 | 38 | 97 | 3 |

### Do methanogens cluster together, despite their phylogenetic diversity?

We identified one module of 128 protein families, module 65 (**Figure 3 and Supplementary Dataset – Table S3)**, that is common to essentially all methanogens, despite the fact that methanogens are not monophyletic **(Figure 1)**. This module contained *mcrA* (Fam05485), a key gene in methane production (Ermler et al. 1997) all the other subunits (BCDG) of methyl–coenzyme M reductase (Mcr), five subunits of the methyl-tetrahydromethanopterin (methyl-H4MPT): coenzyme M methyltransferase (Mtr), five hypothetical conserved proteins in methanogens (Borrel et al. 2014) and genes for transport of iron, magnesium, cobalt and nickel and for synthesis of key cofactors that are required for growth of methanogens. Details are provided in the **Supplementary Materials**.

Modules 72, 129 and 184 are enriched in subunits of the energy-converting hydrogenase A and B and in enzymes for the utilization of methanol (fam04064 and fam05405), methylamine (fam02336 and fam03937), dimethylamine (fam03076 and fam05873), and trimethylamine (fam04092 and fam21299), which are substrates for methanogenesis (Burke, Lo, and Krzycki 1998) (for details, see the **Supplementary Materials**).

Interestingly, we recovered mcr subunits in lineages that are not considered as canonical methanogenic lineages (Evans et al. 2019). These include two genomes of Bathyarchaeota related to BA1 and BA2 (GCA_002509245.1 and GCA_001399805.1) (Evans et al. 2015), and one Archaeoglobi genome related to JdFR-42 (GCA_002010305) (Boyd et al. 2019; Wang et al. 2019). These genomes have been described as having divergent MCR genes. It is reassuring that our method is sensitive enough to recover distant homology. Overall, the correspondence between the distribution of protein families linked to methanogenesis and methanogens supports the validity of our protein family delineation method (**Supplementary Figure 7**).

### Functions specific to Thalassoarchaea

Modules 32 and 71, encompassing 199 families, were consistently associated with genomes of Thalassoarchaea (**Figure 3 and Supplementary Dataset – Table S3)**, which are implicated in protein and saccharide degradation (Tully 2019) (for details, see the **Supplementary Materials**). These modules contain protein degrading enzymes (several different classes of peptidases and one oligotransporter) previously found in Thalassoarchaea (Tully 2019) and two new Thalassoarchaea-specific families of well-conserved peptidases. As reported by (Tully 2019), peptidase S15 (PF02129; fam03321) and peptidase M60-like (PF13402; fam05454) have a narrow distribution within Thalassoarchaea, and were not assigned to ones of the 96 modules. Interestingly, we identified modules specific to Thalassoarchaea subgroup a (MGIIa) (module_135, containing 99 families) and Thalassoarchaea subgroup b (MGIIb) (module_45, containing 39 families) with calcium-binding domains (**Supplementary Figure 8**). These proteins may be involved in signaling and regulation of protein-protein interactions in the cell (Michiels et al. 2002).

### Functions specific to Crenarchaeota

The Crenarchaeota comprises thermophilic organisms that are divided into three main classes, the Thermoproteales, the Sulfolobales and the Desulfurococcales. Two distinct modules with distinct distributions were retrieved. Module 66 (61 families) is widespread in the three classes of Crenarchaeota whereas module 2 (215 families) is specific to the Sulfolobales class **(Figure 3 and Supplementary Dataset – Table S3)**. Interestingly, the subunits of RNA polymerase (Korkhin et al. 2009)), RpoG/Rpb8 (fam03177) are widespread in Crenarchaeota but Rpo13 (fam03159) seems restricted to the *Sulfolobales* class (Korkhin et al. 2009). The Rpo13 protein family of Thermoproteales and Desulfurococcales may be highly divergent from the form described experimentally.

Comparison to PDB enabled annotation of three families with no PFAM and KEGG annotations as having functions related to the DNA replication machinery (**Supplementary Dataset – Table S4)**. We were interested to find that this ubiquitous function is performed by specific protein families in Crenarchaeota, possibly reflecting adaptation to their high temperature habitats. One of these, PolB1-binding protein 2 (PBP2) (fam03141, PDB accession 5n35) (Yan et al. 2017), is a subunit of DNA polymerases B1 (PolB1) that are responsible for initial RNA primer extension with DNA, lagging and leading strand synthesis. The second is a single-stranded DNA-binding protein (DBP) ThermoDBP, which we also found to be conserved in Crenarchaeota and in Thermococci (fam03176, PDB accession 4psl) (Ghalei et al. 2014; Paytubi et al. 2012). Interestingly, however, the third is a Fe-S independent primase subunit PriX (fam03870, PDB accessions: 4wyh and 5of3) specific to Sulfolobales (**Supplementary Figure 9**). PriX is essential for the growth of Sulfolobus cells (Holzer et al. 2017; Liu et al. 2015). These observations point to fundamentally different transcription and replication mechanisms in the major groups within the Crenarchaeota.

Restricted to the Sulfolobales are also two multicopy thermostable acid protease thermopsin families (Lin and Tang 1990) (fam01298 and fam01602 in module 2). Fam01298 is also found in two genomes of Thermoproteales (**Supplementary Figure 9**). Extending a prior report that Crenarchaeota have anomalously large numbers of types I and III CRISPR-Cas systems (Vestergaard, Garrett, and Shah 2014), Crenarchaeota-specific module 66 contains four type I-A Cas families (one of which is the sulfolobales-specific CRISPR-associated protein csaX, fam07252) and four Cas families associated with type III systems (**Supplementary Figure 9**) (**Supplementary Dataset – Table S3)**.

### Functions specific to Thaumarchaeota

The phylum Thaumarcheaota mostly contains aerobic ammonia oxidizing archaea (Brochier-Armanet et al. 2008; Adam et al. 2017). Module 142, which contains 216 families, is specific to Thaumarchaeota (**Figure 3 and Supplementary Dataset – Table S3**). Although this module contains protein families for the three subunits of the ammonia monooxygenase, these three families are absent in genomes for two basal Thaumarcheota lineages, as expected based on prior analyses (Adam et al. 2017) (**Supplementary Figure 10**). This module also contains a highly conserved hypothetical family (fam08021), referred to as AmoX (Bartossek et al. 2012), that is known to co-occur with the amoABC genomic cluster (**Supplementary Dataset – Table S5)**. Importantly, essentially all other protein families in Module 142 currently lack functional annotations (**Supplementary Dataset – Table S3)**.

### Functions specific to Thermococci

The Thermococci comprises sulfur-reducing hyperthermophilic archaea (Palaeococcus, Thermococcus and Pyrococcus). Module 8 contains 146 families abundant in Thermococci and absent in other archaeal lineages (**Figure 3 and Supplementary Dataset – Table S3)**. For example, 98% of the Thermococci genomes have a group 3b (NADP-reducing) [NiFe] hydrogenase. This hydrogenase, also known as sulfhydrogenase, is likely bidirectional (Schut et al. 2012). Only the subunit beta of the sulfur reductase (fam04571) is present in module 8. Subunits alpha (fam00341), delta (fam00630) and gamma (fam00435) are present in the core module (module 1), probably because they are homologs of other hydrogenases. We also detected hydrogen gas-evolving membrane-bound hydrogenases (MBH) in every Thermococci genome (fam03754 in module 8) (Yu et al. 2018; Schut et al. 2016) (**Supplementary Figure 11)**. The MBH transfers electrons from ferredoxin to reduce protons to form H2 gas (Sapra, Bagramyan, and Adams 2003). The Na^+^-translocating unit of the MBH enables H2 gas evolution by MBH to establish a Na^+^ gradient for ATP synthesis near 100 °C in *Pyrococcus furiosus (Yu et al. 2018).* As with the sulfhydrogenase, only the subunit I of the MBH is present in module 8, other subunits of MBH are present in core modules 1 and 23 probably because MBH-type respiratory complexes are evolutionarily and functionally related to the Mrp H+/Na+ antiporter system (Yu et al. 2018).

In the Thermococci-specific module 8 we detected the alpha and gamma subunits (represented by fam10869 and fam02435, respectively) of the Na^+^-pumping methylmalonyl-coenzyme A (CoA) decarboxylase that performs Na^+^ extrusion at the expense of the free energy of decarboxylation reactions (Dimroth 1987; Buckel 2001). The beta and delta subunit, fam02317 and fam00273, are present in the core module 1, again probably because they are homologs of proteins that perform different functions.

Interestingly, three families from module 8 are encoded adjacent in the Thermococci genomes (fam15060, fam07594 and fam05926) (**Supplementary Dataset – Table S6)**. These are annotated as of fungal lactamase (renamed prokaryotic 5-oxoprolinase A, pxpA) and homologs of allophanate hydrolase subunits (renamed pxpB and pxpC) and are likely to form together an 5-oxoprolinase complex (Niehaus et al. 2017). While oxoproline is a major universal metabolite damage product and oxoproline disposal systems are common in all domains of life, the system encoded by these three families appears to be highly conserved in Thermococci (**Supplementary Figure 11)**.

We found the ribosomal protein L41e (fam02171) (Yutin et al. 2012) in 83% of the genomes of Thermococci but sparsely distributed or absent in other lineages. It has previously been noted that the distribution of L41e in Archaea is uncertain (Lecompte et al. 2002).

Using PDB, we established annotations for three families in Thermococci-specific module 8 that lacked PFAM or KEGG annotations (**Supplementary Dataset – Table S4)**. The first appears to be a small protein that inhibits the proliferating cell nuclear antigen by breaking the DNA clamp in *Thermococcus kodakarensis* (fam09868) (Altieri et al. 2016). The second is the S component of an energy-coupling factor (ECF) transporter (fam02033) likely responsible for vitamin uptake (Zhang, Wang, and Shi 2010). The third (fam01133) is the Valosin-containing protein-like ATPase (VAT) that in *Thermoplasma acidophilum* functions in concert with the 20S proteasome by unfolding substrates and passing them on for degradation (Huang et al. 2016). Finally, three peptidases were detected in module 8 (fam01338, fam26972 and fam05052), thus may be specific to the Thermococci (**Supplementary Figure 11)**.

### Functions specific to Halobacteria

We found that 525 families comprise the Halobacteria-specific modules 13 and 108. Module 108 is composed almost completely of hypothetical proteins (**Figure 3 and Supplementary Dataset – Table S3).**

Module 13 contains the two subunits I (fam02395) and II (fam06634) of the high-affinity oxygen cytochrome *bd* oxidase (module 13) and was identified in half of the genomes (**Supplementary Fig. 12)**. It also contains three families without KEGG and PFAM annotations, but close inspection using HMM-HMM comparison showed that they have distant homology with cytochrome-related proteins (**Supplementary Dataset – Table S4)**. The first, fam02696, has distant homology with the catalytic subunit I of heme-copper oxygen reductases (fam00581) and the genes often colocalize with heme-copper oxygen reductases-related genes such as type C *(cbb3)* subunit I or the nitric oxide reductase subunit B (fam00581) **(Supplementary Dataset – Table S7)**. The two other families are cytochrome c associated proteins (fam01001, cytochrome c biogenesis factor and fam02143, cytochrome C and quinol oxidase polypeptide I). Consistent with the presence of oxygen respiration-related families, a catalase-peroxidase gene is present in 90% (fam02210) of the halobacteria genomes (**Supplementary Fig. 12**). Module 13 also includes proteins for synthesis of proteinaceous gas vacuoles (fam03834, fam03740, fam02854 and fam00889; identified in more than 45% of halobacterial genomes, **Supplementary Dataset – Table S3**) that regulate buoyancy of cells in aqueous environments (DasSarma and Arora 2006). The module also includes bacterioruberin 2”, 3”-hydratase (fam00736, CruF; identified in 97% of the halobacteria genomes). Adjacent in the Halobacteria genomes are two families found in the core module 1 (fam00008 and fam00115) and annotated as digeranylgeranylglycerophospholipid reductase and UbiA prenyltransferases respectively **(Supplementary Dataset – Table S7)**. Closer inspection of these three co-encoded enzymes in *Haloarcula japonica* DSM 6131 (GCA_000336635.1) showed they are identical with the bifunctional lycopene elongase and 1,2-hydratase (LyeJ, fam00115) and the carotenoid 3,4-desaturase (CrtD, fam00008) and the bacterioruberin 2”, 3”-hydratase (CruF, fam00736) genes described in *Haloarcula japonica* JCM 7785^T^ (Yang et al. 2015). Together, these three enzymes can generate C50 carotenoid bacterioruberin from lycopene in *Haloarcula japonica (Yang et al. 2015).* Our results showed that C50 carotenoid bacterioruberin is highly conserved in Halobacteria (**Supplementary Figure 12**).

### Functions specific to the six Asgard genomes

The module 48 contains 42 families that are specific and conserved in the six genomes of the superphylum Asgard (four genomes of Thorarchaeota and two genomes of Heimdallarchaeota) (**Figure 3)**. Of these, 33 lack both KEGG and PFAM functional predictions (**Supplementary Dataset – Table S3)**. The Asgard archaea, which affiliate with eukaryotes in the tree of life (Cox et al. 2008), encode many proteins that they share with eukaryotes (Hartman and Fedorov 2002). We detected four eukaryotic signature protein families (ESPs) in module 48 that were described in previous studies (**Supplementary Figure 13 and Supplementary Materials**) (Zaremba-Niedzwiedzka et al. 2017; Spang et al. 2015; Akil and Robinson 2018).

Interestingly, we found a family in module 48 (fam15271) that shows sequence similarity with the integrin beta 4. To the best we know, integrin genes were never described in archaea and fam15271 may represent a new ESP. The genes of fam15271 are always located next to tubulin genes (fam00241) in the five Asgard genomes **(Figure 4A and Supplementary Dataset – Table S8)**. This is particularly interesting as recent studies have shed light on the crosstalk between integrin and the microtubule cytoskeleton (LaFlamme et al. 2018). Finally, one family in module 48 (fam18955) is annotated as the DNA excision repair protein ERCC-3 in three Asgard genomes and three Theionarchaea genomes. The genes neighboring the genes of fam18955 differ between the two lineages (**Figure 4B and Supplementary Dataset – Table S8**) and the three Asgard sequences only share between 20 and 23% protein identity with the three Theionarchaea sequences. These differences may indicate two distinct functions for this family. Fam18955 shows distant homology with the protein RAD25 of *Saccharomyces cerevisiae.* RAD25 is a DNA helicase required for DNA repair and RNA polymerase II transcription in *S. cerevisiae* (Guzder et al. 1994). RAD25 is also one of the six subunits of the transcription factor IIH (TFIIH) in *S. cerevisiae* (Sung et al. 1996). Consistent with the role of RAD25 in *S. cerevisiae,* the genes of family18955 is found next to replication factor C small subunit genes in the three Asgard genomes (**Figure 4 and Supplementary Dataset – Table S8)**.

**Figure 4.**
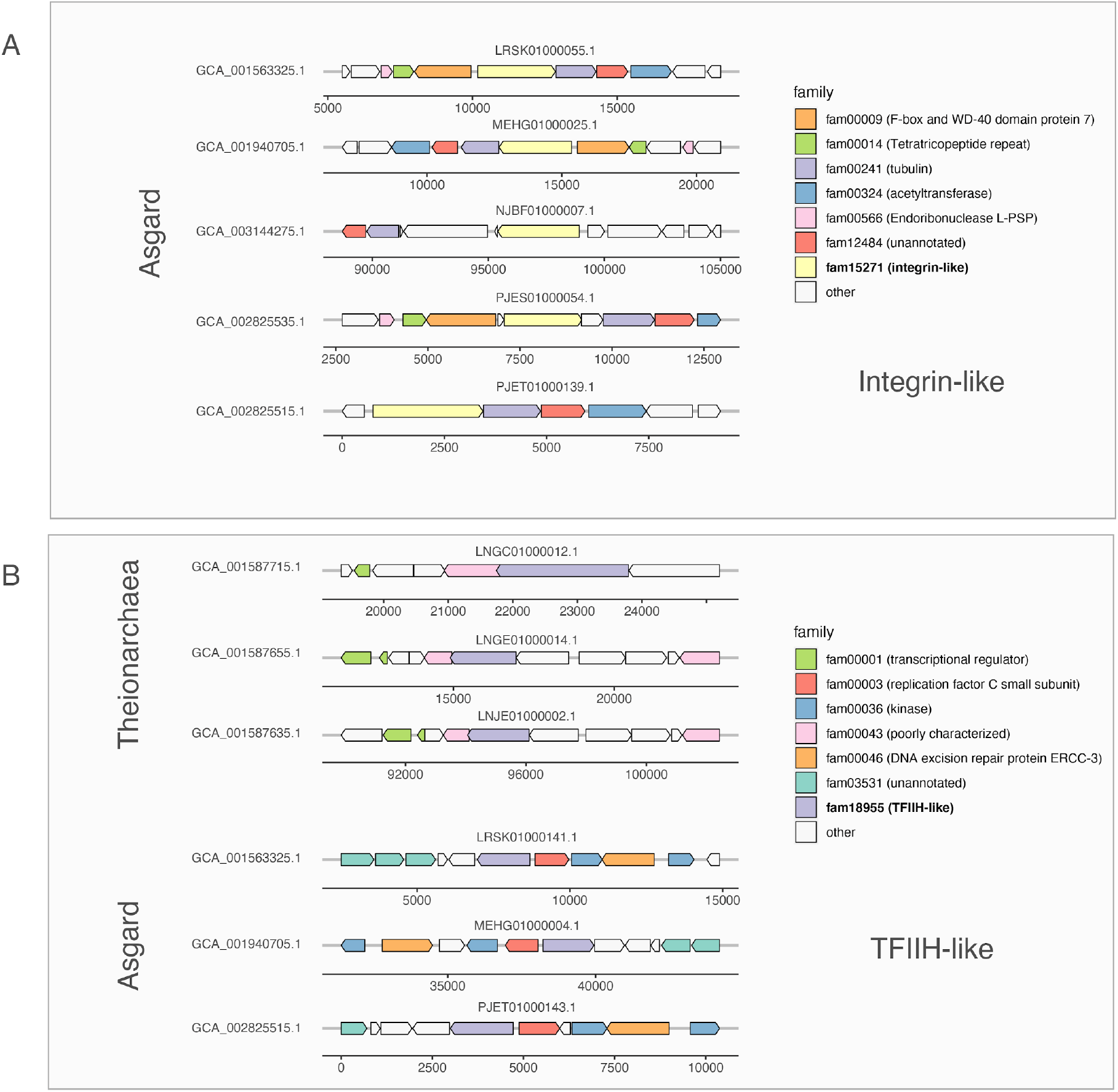
Schematic overview of integrin-like and TFIIH-like gene clusters identified in archaea. A. Conserved gene clusters comprising archaeal integrin-like genes (fam15271) identified in five Asgard genomes. B. Conserved gene clusters comprising archaeal TFIIH-like genes (fam18955) identified in three Theionarchaea and three Asgard genomes. A full gene synteny and genomic context of the genes neighboring the integrin-like (fam15271) and TFIIH-like (fam18955) genes is available in Supplementary Dataset – Table S8.

### Groups without lineage-specific metabolic signatures

The Archaeoglobi and Thermoplasmata lineages are unusual in that they have modules specific to them (modules 105 and 171 respectively), but no specific capacities were identified only in these groups based on functional predictions (**Supplementary Dataset – Table S3**). These lineage-specific modules have the highest percentage of hypothetical families of any lineagespecific module (**Table 1**).

Bathyarchaeota is the only lineage having more than 10 genomes and that does not have a specific module of families that is widespread in the 19 Bathyarchaeota genomes (**Supplementary Dataset – Table S2**). This is intriguing as Bathyarchaeota are widespread across terrestrial marine ecosystems and are known to thrive in diverse chemical environments (Kubo et al. 2012). Alternatively, Bathyarchaeota may be a superphylum. In this case, the lack of modules shared across a superphylum would reinforce the results noted above.

### Hypothetical proteins distinguish the major archaeal groups

Even after augmenting functional predictions using PDB and EggNog databases, families with functional predictions represent a tiny proportion of the protein families that comprise the lineage-specific modules (**Table 1, Figure 5 and Supplementary Dataset – Table S3**). 1053 out of 1411 hypothetical families remain unannotated (**Supplementary Dataset – Table S4**). 358 hypothetical families have small domain matches, but not enough information is available to predict functions. For example, many have domains with matches to zinc finger domains, but such domains occur in proteins with diverse functions (**Supplementary Dataset – Table S4**). We found that the hypothetical proteins are shorter than proteins from the core families of module 1 **(Supplementary Figure 14)** and are more likely to have a transmembrane helix prediction and a signal peptide prediction **(Table 1)**.

Previous studies highlighted the presence of numerous families of proteins with roles in metabolism that are of bacterial origin but occur only in specific archaeal phyla (Nelson-Sathi et al. 2015, 2012). Consequently, we compared all archaeal protein families against a database of bacterial genomes sampled from across the bacterial tree of life to determine the extent to which proteins acquired from bacteria contribute to the archaeal group specific modules (**see materials and methods**). From 3% (Thermoplasmata) to 34% (Halobacteria) of the protein families in modules that are archaeal group specific have homologs in ≥10 distinct bacterial genomes, with the exception of Methanomicrobia, where 63% of the protein families have bacterial homologs (**Table 1**). Thus, for almost all archaeal groups, the majority of the protein families that form modules that separate them from other archaeal groups did not evolve in (or were not acquired from) bacteria. Further, we conclude genes acquired from bacteria only account for a small fraction of the lineage-specific families.

## Discussion

We constructed a set of protein families for Domain Archaea, each of which generally corresponds with a set of homologous proteins with the same predicted function (in cases where functions could be assigned). Protein families with functional predictions that are specific to certain archaeal lineages (e.g., genes involved in methanogenesis or ammonia oxidation) well predict functional traits specific to these lineages. These observations indicate that the protein family construction method is robust. The generated set of 10,866 protein families is provided as an important community research resource. The patterns of presence/absence of protein families across genomes highlight sets of co-occurring proteins (modules), and groupings of genomes based on these modules mostly recapitulate archaeal phylogeny.

What is most striking from our analyses is the prominence of families of hypothetical proteins in the sets of highly prevalent lineage-specific proteins. An important consideration is whether (i) divergence of the sequence of these proteins from proteins with known function simply precluded functional annotation or (ii) whether they are novel proteins that serve well known functions, or if (iii) they represent functions that are unique and evolved following the divergence of each lineage from other archaea. Our analyses were designed to avoid case (i) by relying on state-of-the-art HMM-based homology detection methods that appear to well-group proteins with shared functions (**Supplementary Figure 3**). However, the fact that we could identify some probable functions using protein modeling suggests that (i) is correct in at least a subset of cases. For instance, PriX (fam03870) has structural homology with PriL but no sequence similarity was detected between PriX and any other protein in our analysis. Both proteins are distinct components of the primase complex in *Sulfolobus solfataricus* suggesting that PriX evolved from PriL by duplication followed by subfunctionalization (Holzer et al. 2017; Liu et al. 2015). Lineagespecific hypothetical proteins that are actually homologs of known proteins but currently too divergent for functional assignment are interesting, as they may have been under pressure to evolve more rapidly than normal during lineage divergence. It is not possible to distinguish (ii) from (iii) with the data available. In general, the sets of relatively common archaeal proteins without functional assignments provides targets for future biochemical studies.

Overall, the prevalence of transmembrane helices and signal peptides in the hypothetical proteins in lineage-specific modules indicates that they are membrane associated or extracellular, thus possibly involved in cell-cell and cell-environment interactions (some may be transporters). Where the lineages are confined to specific environments (e.g., halophiles), lineage-specific protein families may have evolved to meet the requirements of that environment (case (i) or (iii)). It is important to note that some modules contain many protein families and probably represent combinations of new functions that, at the present time, cannot be resolved. Regardless of the explanation, or combination of explanations, for the presence of large numbers of lineage-specific proteins, the results clearly indicate the importance of divergence or evolution of a specific subset of proteins during emergence of the major archaeal lineages.

Possibly also informative regarding archaeal evolution is the observation that, despite resolving a Domain-wide core module (module 1), we detected only one case where a clearly defined module is conserved at the superphylum level. It is important to note that, with additional genomes, the two newly recognized Asgard phyla may be reclassified into a single phylum, eliminating this exception. The apparent lack of protein family module support for superphyla may argue against the phyletic gradualism, in which one lineage gradually transforms into another, and favor the theory of cladogenesis, where a lineage splits into two distinct lineages (Gould and Eldredge 1977). We acknowledge that modules containing fewer than 20 protein families (the cutoff used to define modules) may be uniquely associated with superphyla, and that some potentially important archaeal lineages were not included in the current analysis due to lack of a sufficient number of high-quality genomes.

The observation that the patterns of presence/absence of shared protein families groups together archaea that historically have been assigned to the same lineage and separates them from other lineages, in combination with innumerable prior publications on archaeal physiology and taxonomy (Adam et al. 2017; Spang, Caceres, and Ettema 2017; Baker et al. 2020), supports the value of the current taxonomic classifications within Domain Archaea. Overall, the results reinforce the concept that early archaeal evolution rapidly generated the major lineages, the rise of which was linked to acquisition of a set of proteins (recognized here as modules) that were largely retained during subsequent evolution of each lineage.

## Methods

### Genome collection

569 unpublished genomes were added to the 2,618 genomes of Archaea downloaded from the NCBI genome database in September 2018.

132 genomes were obtained from metagenomes of sediment samples. Sediment samples were collected from the Guaymas Basin (27°N0.388, 111°W24.560, Gulf of California, Mexico) during three cruises at a depth of approximately 2000 m below the water surface. Sediment cores were collected during two Alvin dives, 4486 and 4573 in 2008 and 2009. Sites referred to as “Megamat” (genomes starting with “Meg”) and “Aceto Balsamico” (genomes starting with “AB” in name), Core sections between 0-18 cm from 4486 and from 0-33 cm 4573 and were processed for these analyses. Intact sediment cores were subsampled under N_2_ gas, and immediately frozen at −80 °C on board. The background of sampling sites was described previously (Teske et al. 2016). Samples were processed for DNA isolation from using the MoBio PowerMax soil kit (Qiagen) following the manufacturer’s protocol. Illumina library preparation and sequencing were performed using Hiseq 4000 at Michigan State University. Paired-end reads were interleaved using interleav_fasta.py (https://github.com/jorvis/biocode/blob/master/fasta/interleave_fasta.py) and the interleaved sequences were trimmed using Sickle (https://github.com/najoshi/sickle) with the default settings. Metagenomic reads from each subsample were individually assembled using IDBA-UD with the following parameters: --pre_correction --mink 65 --maxk 115 --step 10 -- seed_kmer 55 (Peng et al. 2012). Metagenomic binning was performed on contigs with a minimum length of 2000 bp in individual assemblies using the binning tools MetaBAT (Kang et al. 2015) and CONCOCT (Alneberg et al. 2014), and resulting bins were combined with using DAS Tool (Sieber et al., n.d.). CheckM lineage_wf (v1.0.5) (Parks et al. 2015) was used to estimate the percentage of completeness and contamination of bins. Genomes with more than 50% completeness and 10% contamination were manually optimized based on GC content, sequence depth and coverage using mmgenome (Karst, Kirkegaard, and Albertsen, n.d.).

The remaining 447 genomes came from previous sequencing and binning efforts (genomes starting with “ggkbase”). In brief, 168 genomes were obtained from an aquifer adjacent to the Colorado River near the town of Rifle, Colorado, USA in 2011 (Anantharaman et al. 2016), 50 genomes from the Crystal Geyser system in Utah, USA (Probst et al. 2014). For DNA processing and sequencing methods see (Probst et al. 2017; Anantharaman et al. 2016). Forty-one genomes were obtained from (Parks et al. 2017). Additionally, 188 genomes were obtained from groundwater samples from Genasci Dairy Farm, located in Modesto, California (CA) as described in (He et al., n.d.).

### Genome completeness assessment and de-replication

Genome completeness and contamination were estimated based on the presence of single-copy genes (SCGs) as described in (Olm et al. 2017; Anantharaman et al. 2016). Genome completeness was estimated using 38 SCGs. For non-DPANN archaea, genomes with more than 26 SCGs (>70% completeness) and less than 4 duplicated copies of the SCGs (<10% contamination) were considered as draft-quality genomes. Due to the reduced nature of DPANN genomes (Castelle et al. 2015), genomes with more than 22 SCGs and less than 4 duplicated copies of the SCGs were considered as draft-quality genomes. Genomes were de-replicated using dRep (Olm et al. 2017) (version v2.0.5 with ANI > 95%). The most complete and less contaminated genome per cluster was used in downstream analyses.

### Concatenated 14 ribosomal proteins phylogeny

A maximum-likelihood tree was calculated based on the concatenation of 14 ribosomal proteins (L2, L3, L4, L5, L6, L14, L15, L18, L22, L24, S3, S8, S17, and S19). Homologous protein sequences were aligned using MAFFT (version 7.390) (--auto option) (Katoh and Standley 2016), and alignments refined to remove gapped regions using Trimal (version 1.4.22) (--gappyout option) (Capella-Gutiérrez, Silla-Martínez, and Gabaldón 2009). The protein alignments were concatenated, with a final alignment of 1,179 genomes and 2,388 positions. The tree was inferred using RAxML (Stamatakis 2014) (version 8.2.10) (as implemented on the CIPRES web server (Miller, Pfeiffer, and Schwartz 2010)), under the LG plus gamma model of evolution, and with the number of bootstraps automatically determined via the MRE-based bootstopping criterion. A total of 108 bootstrap replicates were conducted, from which 100 were randomly sampled to determine support values.

### Protein clustering

Protein clustering into families was achieved using a two-step procedure as previously described in (Méheust et al. 2019). A first protein clustering was done using the fast and sensitive protein sequence searching software MMseqs2 (version 9f493f538d28b1412a2d124614e9d6ee27a55f45) (Steinegger and Söding 2017). An all-vs-all search was performed using e-value: 0.001, sensitivity: 7.5 and cover: 0.5. A sequence similarity network was built based on the pairwise similarities and the greedy set cover algorithm from MMseqs2 was performed to define protein subclusters. The resulting subclusters were defined as subfamilies. In order to test for distant homology, we grouped subfamilies into protein families using an HMM-HMM comparison procedure as follows: the proteins of each subfamily with at least two protein members were aligned using the result2msa parameter of mmseqs2, and, from the multiple sequence alignments, HMM profiles were built using the HHpred suite (version 3.0.3) (Soding 2005). The subfamilies were then compared to each other using hhblits (Remmert et al. 2011) from the HHpred suite (with parameters -v 0 -p 50 -z 4 -Z 32000 -B 0 -b 0). For subfamilies with probability scores of ≥ 95% and coverage ≥ 0.50, a similarity score (probability × coverage) was used as weights of the input network in the final clustering using the Markov Clustering algorithm (Enright, Van Dongen, and Ouzounis 2002), with 2.0 as the inflation parameter. These clusters were defined as the protein families.

### Module definition and taxonomic assignment

Looking at the distribution of the protein families across the genomes, a clear modular organization emerged. Modules of families were defined using a cutoff of 0.95 on the dendrogram tree of the families. The dendrogram tree was obtained from a hierarchical clustering using the Jaccard distance that was calculated based on profiles of protein family presence/absence. The corresponding clusters define the modules.

A phyla distribution was assigned to each module using the method of (Méheust et al. 2019). For each module, the median number of genomes per family (m) was calculated. The genomes were ranked by the number of families they carry. The m genomes that carry the most of families were retained; their phyla distribution defines the taxonomic assignment of the module. Functional annotation Protein sequences were functionally annotated based on the accession of their best Hmmsearch match (version 3.1) (E-value cut-off 0.001) (Eddy 1998) against an HMM database constructed based on ortholog groups defined by the KEGG (Kanehisa et al. 2016) (downloaded on June 10, 2015). Domains were predicted using the same hmmsearch procedure against the Pfam database (version 31.0) (Punta et al. 2012). The domain architecture of each protein sequence was predicted using the DAMA software (version 1.0) (default parameters) (Bernardes et al. 2016). SIGNALP (version 5.0) (parameters: -format short -org arch) (Almagro Armenteros et al. 2019) was used to predict the putative cellular localization of the proteins. Prediction of transmembrane helices in proteins was performed using TMHMM (version 2.0) (default parameters) (Krogh et al. 2001). Protein sequences were also functionally annotated based on the accession of their best hmmsearch match (version 3.1) (E-value cut-off 1e-10) against the PDB database (Rose et al. 2017) (downloaded in February 2020).

### HMM-HMM Predictions

Subfamilies were used to perform HMM-HMM annotation against arCogs of EggNog (version 5.0) (Huerta-Cepas et al. 2019) using hhblits (Remmert et al. 2011) from the HHpred suite (with parameters -v 0 -p 50 -z 4 -Z 32000 -B 0 -b 0). Subfamilies were subsequently functionally annotated based on the EggNog accessions of their best probability score.

### Sequence similarity analysis

The 75,737 subfamilies from the 10,866 families were searched against a bacterial database of 2,552 bacterial genomes (Méheust et al. 2019) using hmmsearch (version 3.1) (E-value cut-off 0.001) (Eddy 1998). Among them 46,261 subfamilies, comprising 8,300 families, have at least one hit against a bacterial genome.

## Data availability

The newly reconstructed genomes have been deposited at NCBI under BioProject **XX** (**TBA**). The genomes of the herein analysed archaea have been made publicly available on the ggkbase database (**TBA**). Detailed annotations of the families are provided in the **Supplementary Dataset – Table S3** accompanying this paper. Raw data files (phylogenetic tree and fasta sequences of the families) are made available via figshare under the following link: **TBA**.

## Author contributions

R.M., C.J.C. and J.F.B. designed the study. R.M. and C.J.C. created the dataset. C.J.C performed the phylogenetic analysis. A.L.J. performed the bacterial analysis. R.M. performed the protein family, the module detection, the genome annotations and HMMs analyses. R.M., C.J.C. and J.F.B. wrote the manuscript. All authors read and approved the final manuscript.

## Materials and Correspondence

Correspondence and material requests should be addressed to.

## Acknowledgements

We thank Dr. Brett Baker, Dr Kiley Seitz and Dr Xianzhe Gong for the permission to use the metagenomic datasets from the Guaymas Basin in this study. We thank Dr. Christine He for the permission to use the metagenomic dataset from Genasci. We acknowledge funding support from the Chan Zuckerberg Biohub and the Innovative Genomics Institute at UC Berkeley.

