## SupplementaryMaterials for "Early acquisition of conserved, lineage-specific proteins currently lacking functional predictions were central to the rise and diversification of archaea"

**for**

**Do methanogens cluster together, despite their phylogenetic diversity? (Full analysis)**

To test whether we could detect expected patterns of protein family distribution across lineages using our approach, we examined the distribution of genes involved in methanogenesis, including the alpha subunit *mcrA*, a key gene in methane production (Ermler et al. 1997). The mcrA protein family (fam05485) was identified in module 65, which comprises 128 protein families and is highly enriched in Methanomicrobia, Methanobacteria and Methanomassiliicoccales (**Supplementary Figure 7**). Along with the *mcrA* gene, all the other subunits (BCDG) of methyl–coenzyme M reductase (Mcr) were identified in module 65 (fam02993, fam03716, fam15638 and fam04416). Five subunits of the methyl-tetrahydromethanopterin (methyl-H4MPT): coenzyme M methyltransferase (Mtr) (BCDEF) were also found (fam06037, fam06163, fam06462, fam06013 and fam02119 respectively) although the *Mtr* genes were absent in *Methanomassiliicoccales* (Borrel et al. 2014) (**Supplementary Dataset - Table S3**). Using HMM-HMM comparison method against the eggNOG database (Huerta-Cepas et al. 2019) (**see materials and methods**), we also detected five hypothetical conserved protein families that are associated with the core proteins of methanogenesis (fam02372, fam05394, fam06062, fam06600 and fam12037) (Borrel et al. 2014) (**Supplementary Dataset - Table S4**). The occurrence between the Mcr subunits and the five methanogenesis markers of unknown functions is striking (**Supplementary Figure 7)**. Further, genes for transport of iron, magnesium, cobalt and nickel and for synthesis of key cofactors that are required for methanogens growth were also found in the module 65. We identified two other modules enriched in subunits of the energy-converting hydrogenase A (fam10726, fam14360, fam17633, fam06266, fam13666, fam08995, fam03063, fam02679, fam22457 and fam06367) and B (fam06098, fam06875 and fam32156) (module 129) and in enzymes for the utilization of methylamine (fam02336 and fam03937), dimethylamine (fam03076 and fam05873), and trimethylamine (fam04092 and fam21299) as substrates for methanogenesis (Burke et al. 1998) (module 184). Methylamine, dimethylamine and trimethylamine are in two distinct families instead of one single family due to mispredictions of the coding sequences (each gene was split in two consecutive genes by the gene prediction software Prodigal). We found two families of methanol---5-hydroxybenzimidazolylcobamide Co-methyltransferase in two distinct families (fam05405 and fam04064) in modules 184 and 72 that are specific to the Methanomassiliicoccales (**Supplementary Figure 7 and Supplementary Dataset - Table S3**). When the distribution of the signature families is rendered on the phylogenetic tree of Archaea the correspondence between families, modules and annotations is apparent (**Supplementary Figure 7**).

We also recovered mcr subunits in lineages that are not considered as canonical methanogenic lineages (Evans et al. 2019). These include two genomes of Bathyarchaeota related to BA1 and BA2 (GCA_002509245.1 and GCA_001399805.1) (Evans et al. 2015), and one Archaeoglobi genome related to JdFR-42 (GCA_002010305) (Boyd et al. 2019; Wang et al. 2019). These genomes have been described as having divergent MCR genes. It is reassuring that our method is sensitive enough to recover distant homology (**Supplementary Figure 7**).

**Functions specific to Thalassoarchaea (Full analysis)**

Modules 32 and 71, encompassing 199 families, were consistently associated with genomes of Thalassoarchaea archaea. Recent comparative genomic analysis of 250 Thalassoarchaea genomes revealed the ecological roles of these archaea in protein and saccharide degradation (Tully 2019). Protein degrading enzymes (several different classes of peptidases and one oligotransporter) found in modules 32 and 71, include some previously linked to Thalassoarchaea archaea (fam05120, fam05272 and fam01092) (Tully 2019). We also identified two new Thalassoarchaea-specific families of well-conserved peptidases that are seemingly unique to Thalassoarchaea (peptidase M17; PF00883; fam03211 (Module 32) and peptidase M6; PF05547 fam00840 (Module 71)). As reported by Tully (2019), peptidase S15 (PF02129; fam03321) and peptidase M60-like (PF13402; fam05454) have a narrow distribution within Thalassoarchaea so were assigned to another module (module_738). Interestingly, we identified modules specific to Thalassoarchaea subgroup IIa (module_135, containing 99 families) and Thalassoarchaea subgroup IIb (module_45, containing 39 families). Both modules contain 4 protein families with calcium-binding domains (fam02857, fam00852 and fam02101 in module 135 and fam01838 in module 45) (**Supplementary Figure 8**). These proteins may be involved in signaling and regulation of protein-protein interactions in the cell (Michiels et al. 2002).

**Functions specific to the six Asgard genomes (Full analysis).**

The module 48 contains 42 families that are specific and conserved in the six genomes of the superphylum Asgard (four genomes of Thorarchaeota and two genomes of Heimdallarchaeota). Of these, 33 lack both KEGG and PFAM functional predictions (**Supplementary Dataset - Table S3**). The Asgard archaea, which affiliate with eukaryotes in the tree of life (Cox et al. 2008), encode proteins that they only share with eukaryotes (Hartman and Fedorov 2002). We detected six eukaryotic signature protein families (ESPs) in module 48 (**Supplementary Figure 13**). These ESPs include the ESCRT-III Snf7-domain family (fam04979) and the ESCRT-II Vps25-like family (fam12378). These two families are part of the ESCRT system and the genes are found in the same genomic region in the Asgard genomes (Zaremba-Niedzwiedzka et al. 2017).

Three cytoskeleton-related families were found in module 48. The two first families are the gelsonin-domain protein family (fam03231) (Zaremba-Niedzwiedzka et al. 2017) and fam04420 that matches with the PDB sequence of the Loki profilin-1 crystal structure (accession: 5yee) (Akıl and Robinson 2018). Interestingly, the third cytoskeleton-related family (fam15271) shows similarity with the integrin beta 4. To the best we know, integrin genes were never described in archaea and fam15271 may represent a new ESP. The genes of fam15271 are always located next to tubulin genes (fam00241) in the five Asgard genomes (**Figure 4A and Supplementary Dataset - Table S8**). This is particularly interesting as recent studies have shed light on the crosstalk between integrin and the microtubule cytoskeleton (LaFlamme et al. 2018).

Finally, one family in module 48 (fam18955) is annotated as the DNA excision repair protein ERCC-3 in three Asgard genomes and three Theionarchaea genomes. The genes neighboring the genes of fam18955 differ between the two lineages (**Figure 4B and Supplementary Dataset - Table S8**) and the three Asgard sequences only share between 20 and 23% protein identity with the three Theionarchaea sequences. These differences may indicate two distinct functions for this family. Fam18955 shows distant homology with the protein RAD25 of Saccharomyces cerevisiae. RAD25 is a DNA helicase required for DNA repair and RNA polymerase II transcription in S. cerevisiae (Guzder et al. 1994). RAD25 is also one of the six subunits of the transcription factor IIH (TFIIH) in S. cerevisiae (Sung et al. 1996). Consistent with the role of RAD25 in S. cerevisiae, the genes of family18955 is found next to replication factor C small subunit genes in the three Asgard genomes (**Figure 4 and Supplementary Dataset - Table S8**).

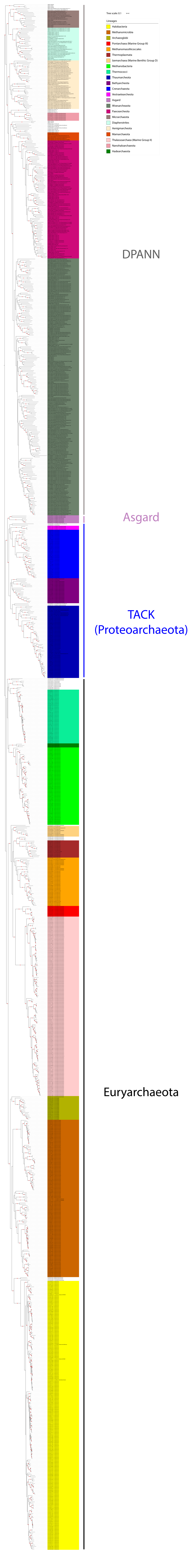

Tree scale: 0.1
Lineages
Halobacteria
Methanomicrobia
Archaeoglobi
Pontarchaea (Marine Group III)
Methanomassiliicoccales
Thermoplasmata
Izemarchaea (Marine Benthic Group D)
Methanobacteria
Thermococci
Thaumarcheota
Bathyarcheota
Crenarchaeota
Vestraetearcheota
Asgard
Woesarcheaota
Paecearcheota
Micrarchaeota
Diapherotrites
Aenigmarcheota
Mamarchaeota
Thalassoarchaea (Marine Group II)
Nanohaloarchaeota
Hadearchaeota
DPANN
Asgard
TACK
(Proteoarchaeota)
Euryarchaeota

**Supplementary Figure S1.** Taxonomic assessment and distribution of the 1,179 representative genomes. Maximum-likelihood phylogeny based on a 14-ribosomal-protein concatenated alignment using the LG plus gamma model of evolution. Scale bar indicates the average substitutions per site.

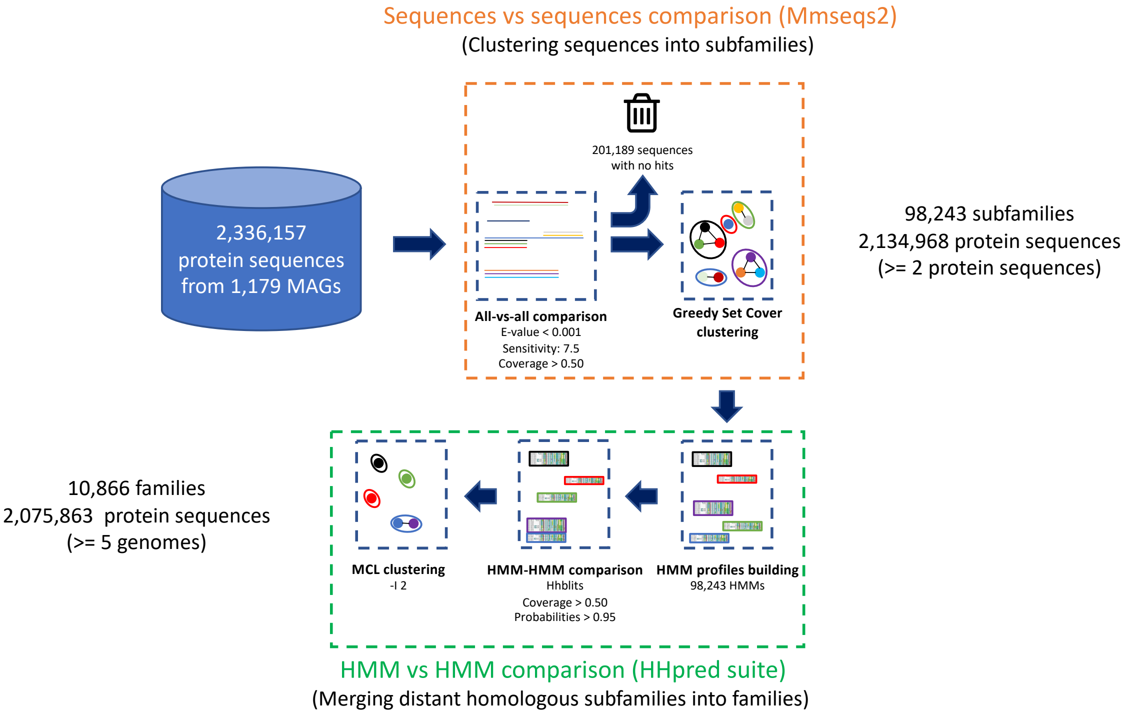


**Supplementary Figure S2.** The protein clustering pipeline used in the study. MAGs: metagenome-assembled genomes.

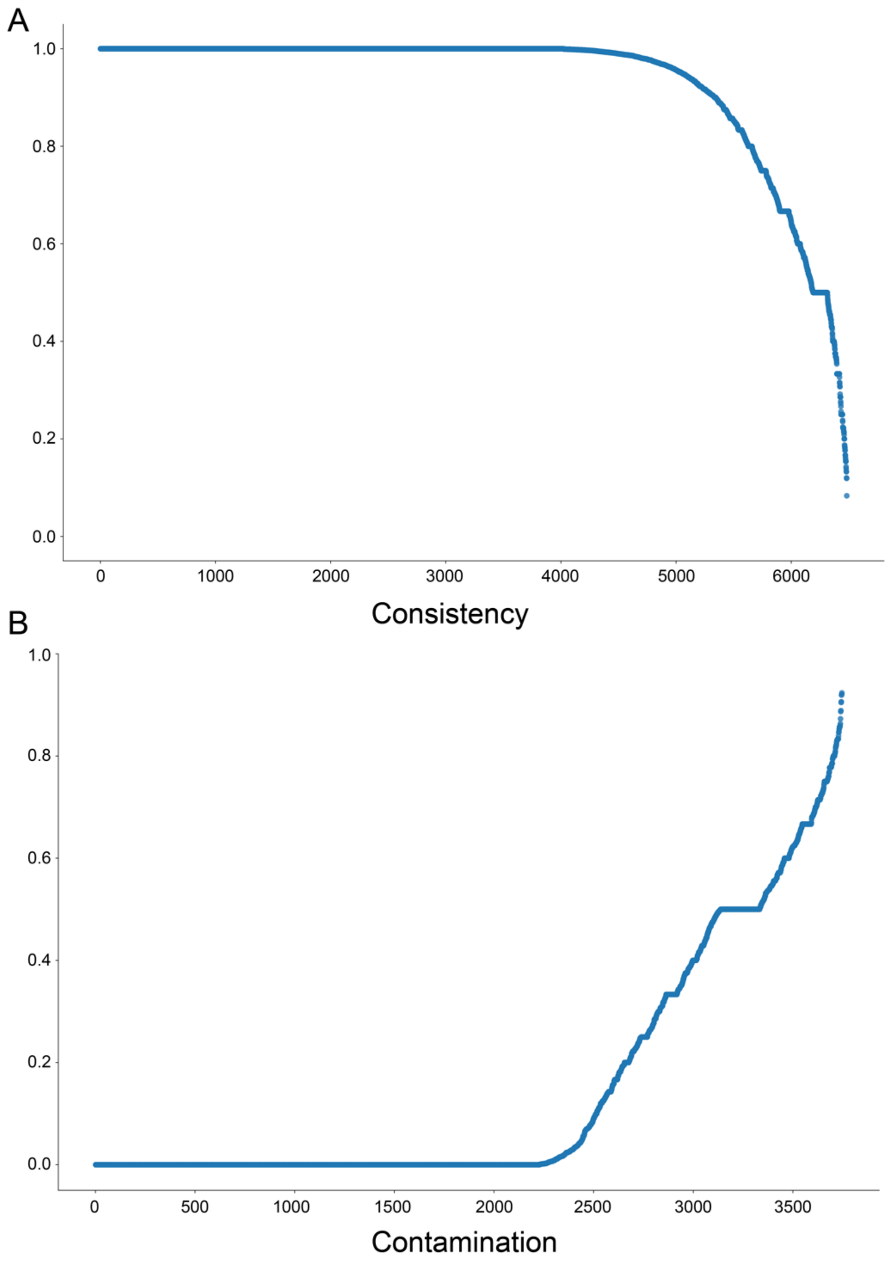


**Supplementary Figure S3.** Quality assessment of the protein clustering. A. Consistency between the KEGG annotations and the protein families. For each of the 6482 annotations, we reported the family which contains the highest percentage of protein members annotated with that KEGG annotation. Each dot represents a KEGG annotation, the y-axis represents the highest percentage. C. Contamination of the protein families. For each family with proteins having KEGG annotations, we computed the percentage of the proteins that have KEGG annotations different than the most abundant one, this percentage defined the annotation admixture (y-axis). Each dot represents a protein family.

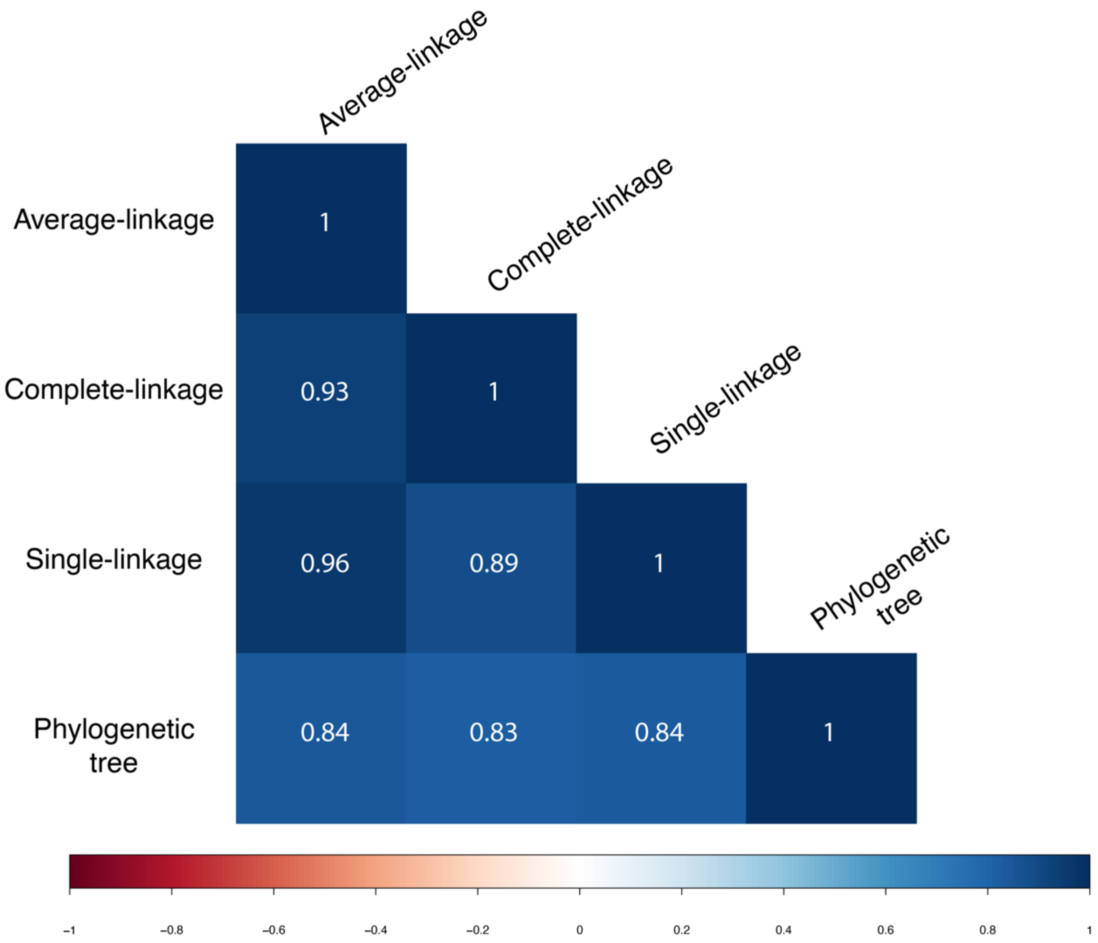


**Supplementary Figure S4.** Correlation plot of 4 trees obtained from 3 different hierarchical clustering methods (complete linkage, average linkage and single linkage). Maximum-likelihood tree based on RAxML is also shown ("Phylogenetic tree"). Correlations are based on cophenetic distance matrices between pairs of trees. Positive correlations are displayed in blue and negative correlations in red color. Color intensity is proportional to the correlation coefficient.

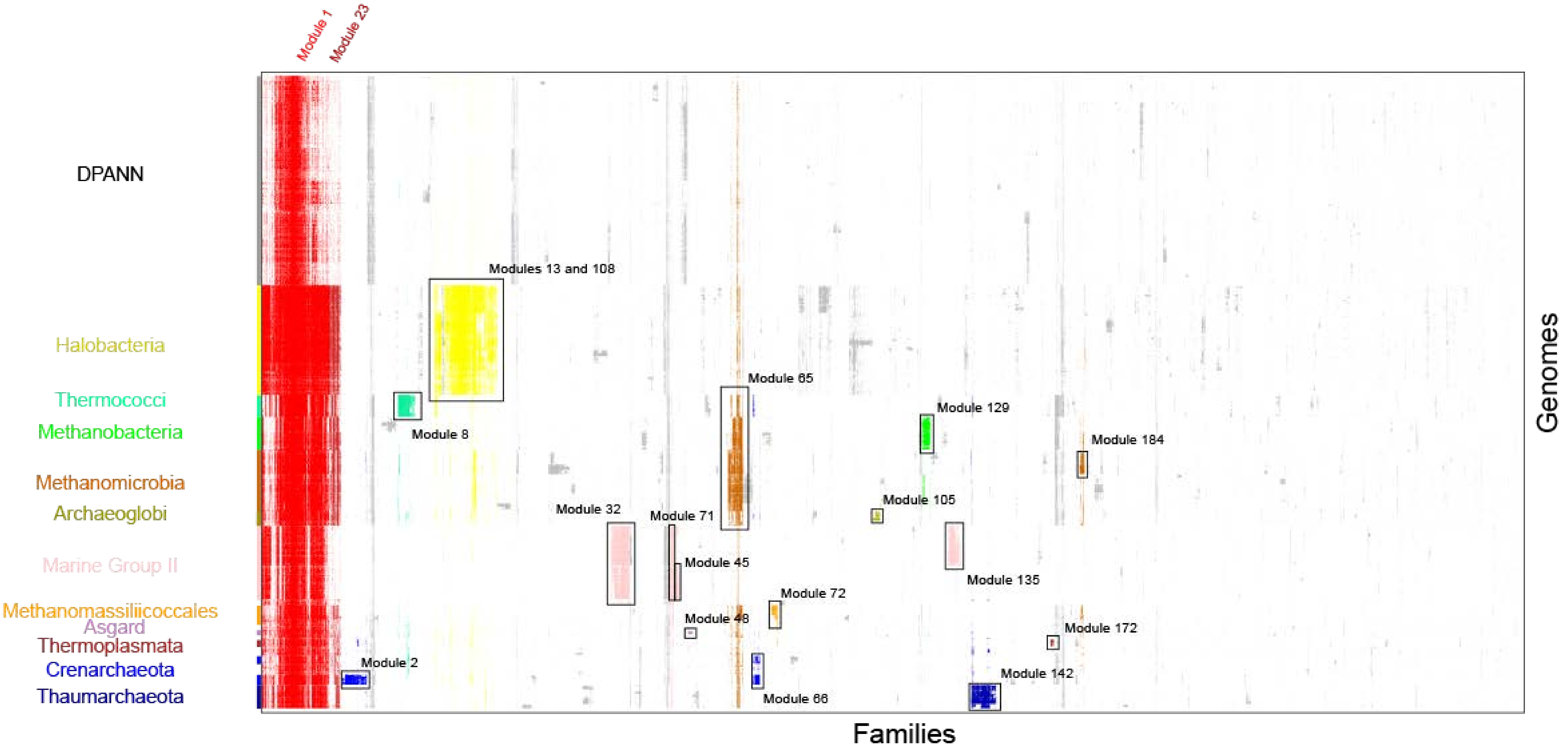

Module 1
Module 23
DPANN
Halobacteria
Thermococci
Methanobacteria
Methanomicrobia
Archaeoglobi
Marine Group II
Methanomassiliicoccales
Asgard
Thermoplasmata
Crenarchaeota
Thaumarchaeota
Modules 13 and 108
Module 8
Module 65
Module 129
Module 184
Module 105
Module 32
Module 71
Module 45
Module 135
Module 72
Module 48
Module 172
Module 2
Module 142
Module 66
Families
Genomes

**Supplementary Figure S5.** The distribution of 10,866 widely distributed protein families (columns) in 1,179 representative genomes (rows) from Archaea. The families of the 19 modules discussed in the study were colored. Data are clustered based on the presence (black) and absence (white) profiles (Jaccard distance, complete linkage).

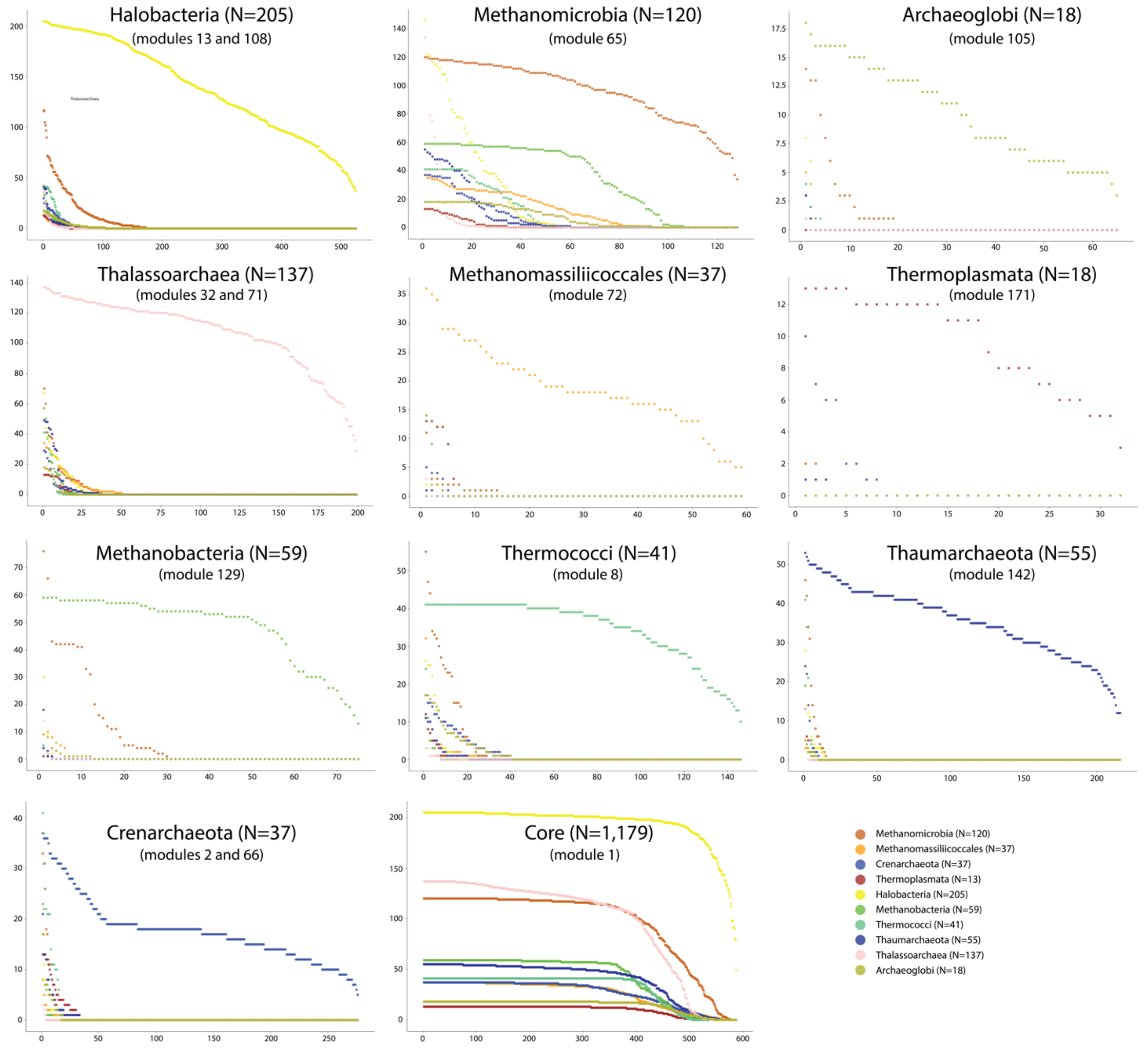


**Supplementary Figure S6.** Number of genomes per family in 14 selected modules. X-axis represents the families and y-axis the number of genomes. For each family, the number of genomes of Methanomicrobia, Methanobacteria, Halobacteria, Crenarchaeota, Thalassoarchaea, Thermococci, Archaeoglobi, Thaumarchaeota, Thermoplasmata and Methanomassiliicoccales is shown by a colored dot.

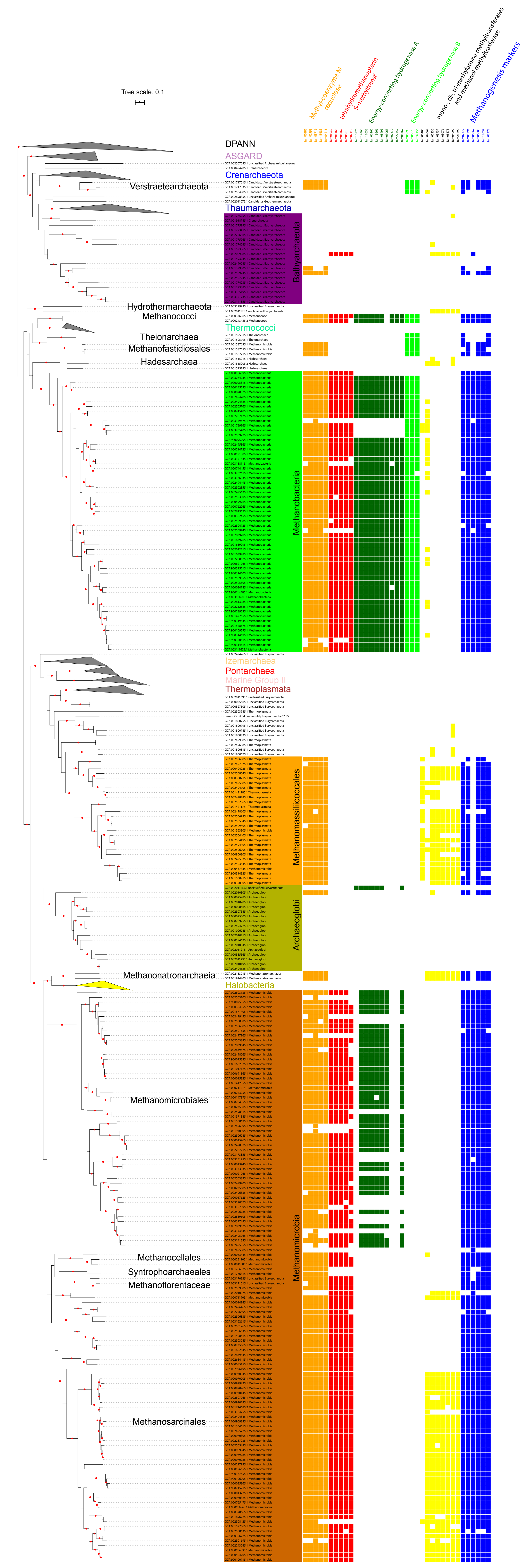
Tree scale: 0.1
Methyl-coenzyme M reductase
tetrahydromethanopterin S-methyltransf
Energy-converting hydrogenase A
Energy-converting hydrogenase B
mono-, di-, tri-methylamine methyltransferases and methanol methyltransferase
Methanogenesis markers
DPANN
ASGARD
Crenarchaeota
Verstraetearchaeota
Thaumarchaeota
Bathyarchaeota
Hydrothermarchaeota
Methanococci
Thermococci
Theionarchaea
Methanofastidiosales
Hadesarchaea
Methanobacteria
Izemarchaea
Pontarchaea
Marine Group II
Thermoplasmata
Methanomassiliicoccales
Archaeoglobi
Methanonatronarchaeia
Halobacteria
Methanomicrobiales
Methanomicrobia
Methanocellales
Syntrophoarchaeales
Methanoflorentaceae
Methanosarcinales

**Supplementary Figure S7.** Presence and absence of 37 families of modules 65, 72, 129 and 184 in genomes of methanogen archaea. Scale bar indicates the average substitutions per site.

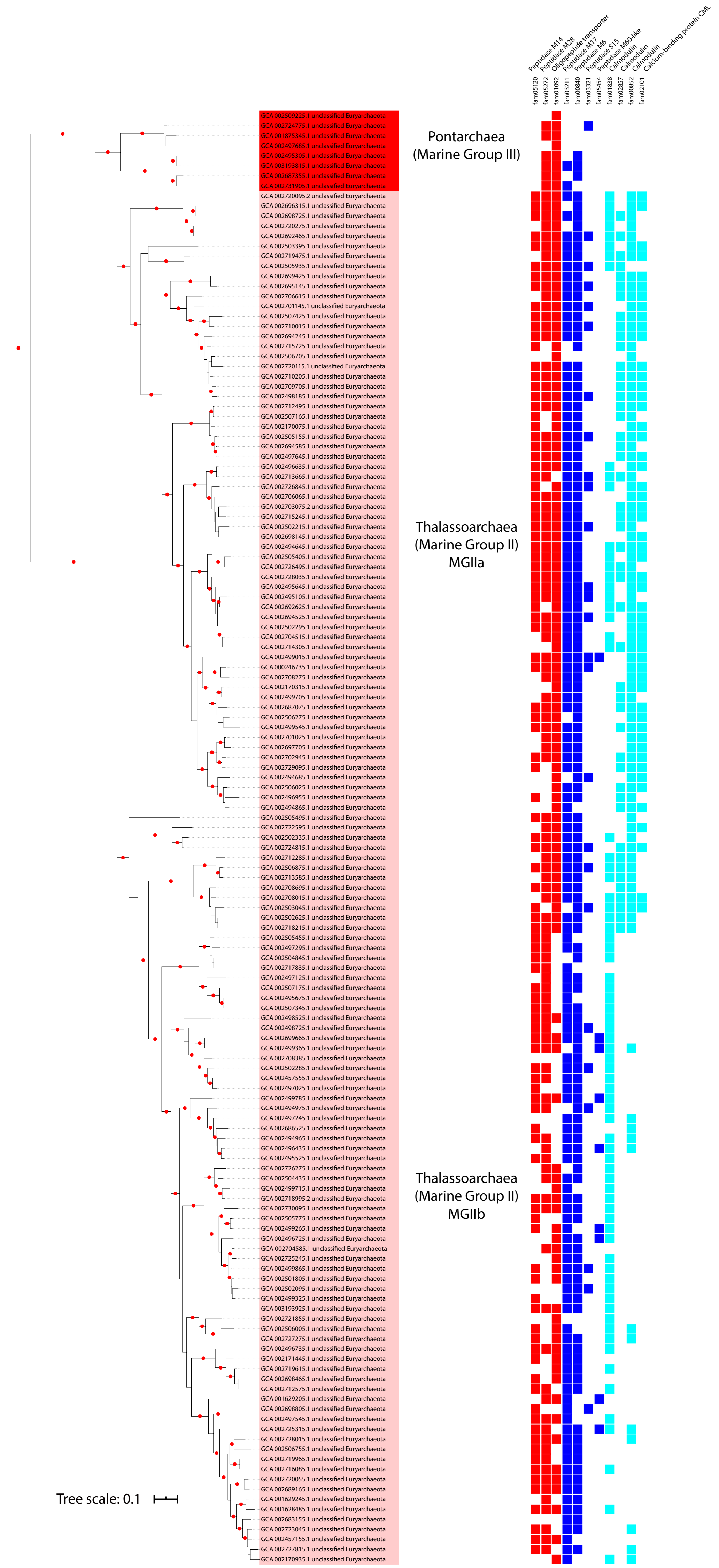

Peptidase M14
Peptidase M28
Oligopeptide transporter
Peptidase M17
Peptidase M6
Peptidase S15
Peptidase M60-like
Calmodulin
Calmodulin
Calcium-binding protein CML
fam05120
fam05272
fam01092
fam03211
fam00840
fam03321
fam05454
fam01838
fam02857
fam00852
fam02101
Pontarchaea
(Marine Group III)
Thalassoarchaea
(Marine Group II)
MGIIa
Thalassoarchaea
(Marine Group II)
MGIIb
Tree scale: 0.1

**Supplementary Figure S8.** Presence and absence of 11 families of modules 32, 45, 71, 135 in the genomes of Thalassoarchaea. Scale bar indicates the average substitutions per site.

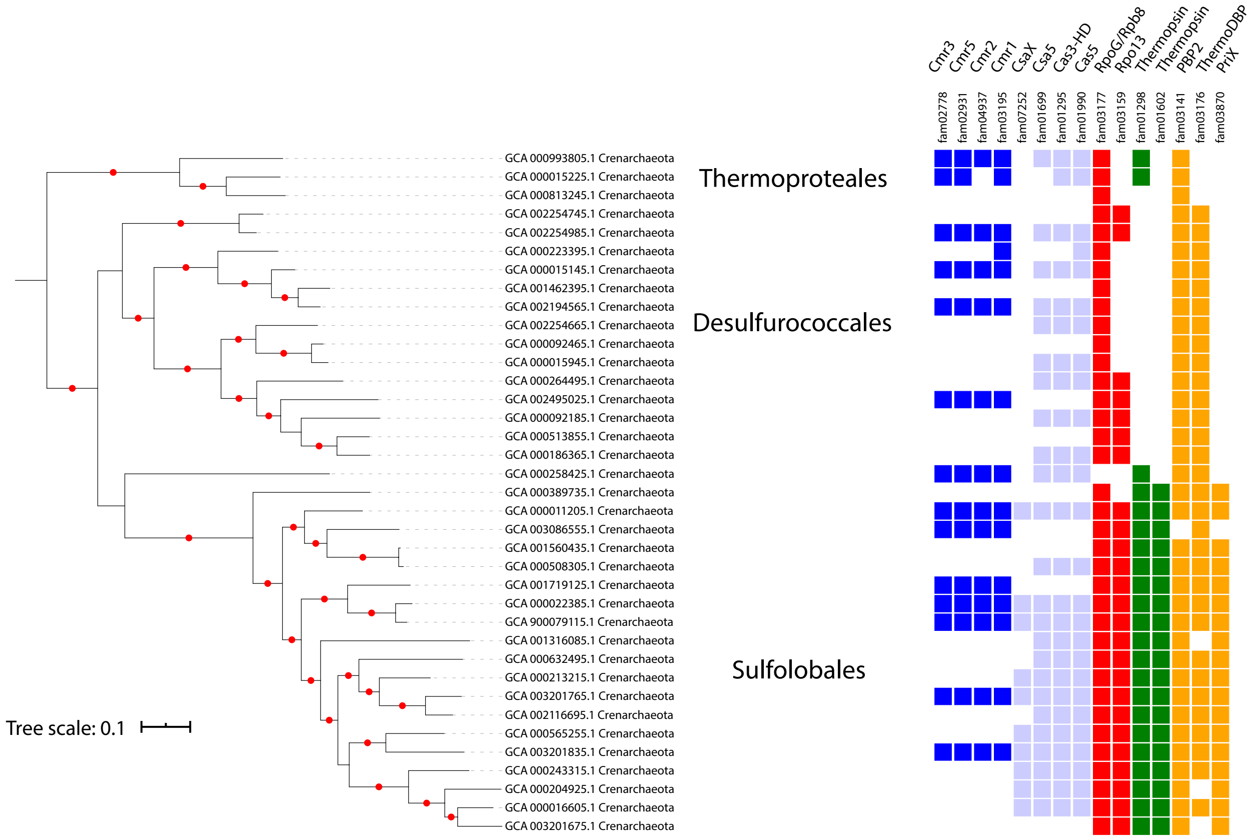


**Supplementary Figure S9.** Presence and absence of 15 families of modules 2 and 66 in genomes of Crenarchaeota. Scale bar indicates the average substitutions per site.

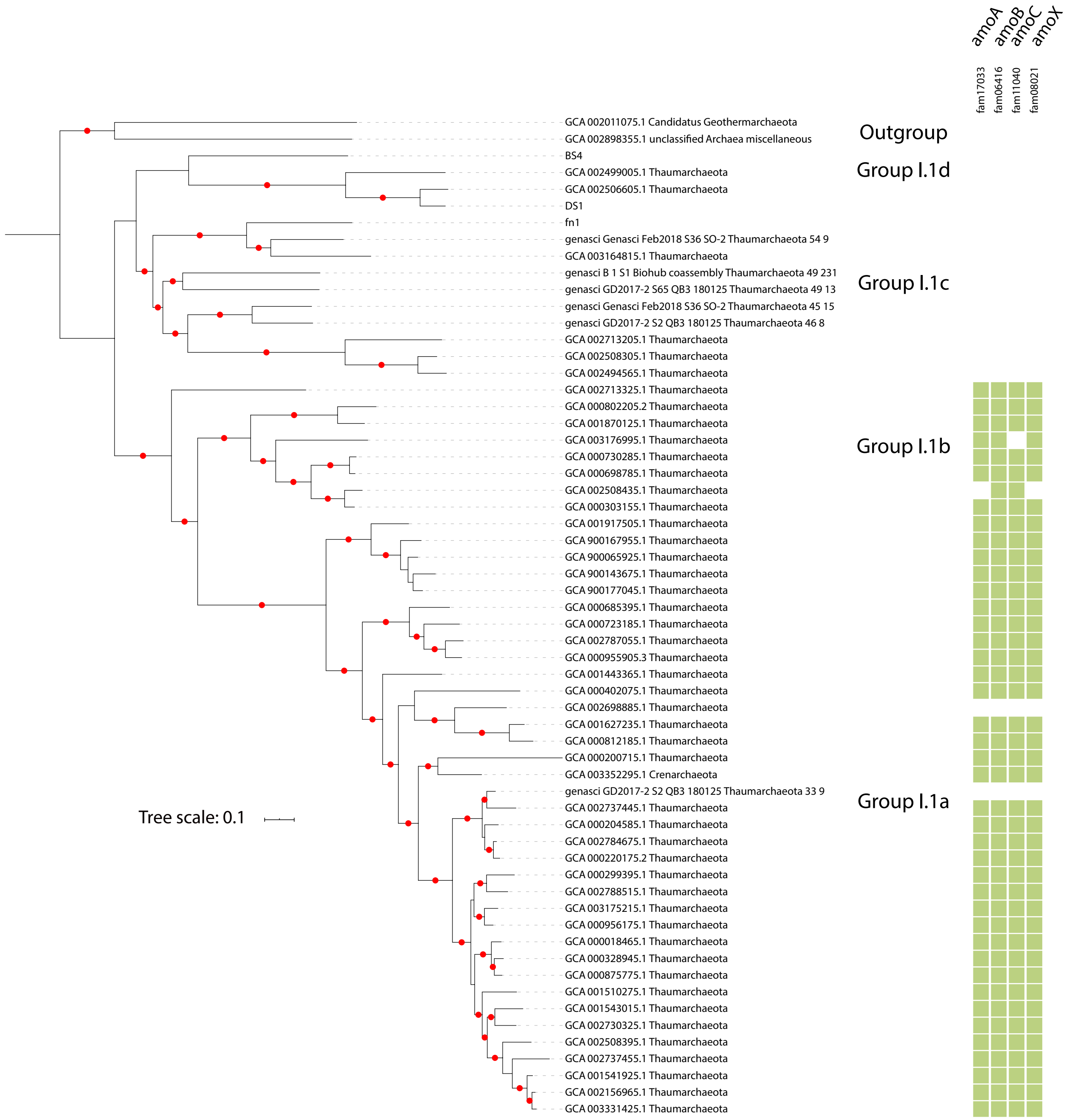


**Supplementary Figure S10.** Presence and absence of 4 families of module 142 in genomes of Thaumarchaeota. Scale bar indicates the average substitutions per site.

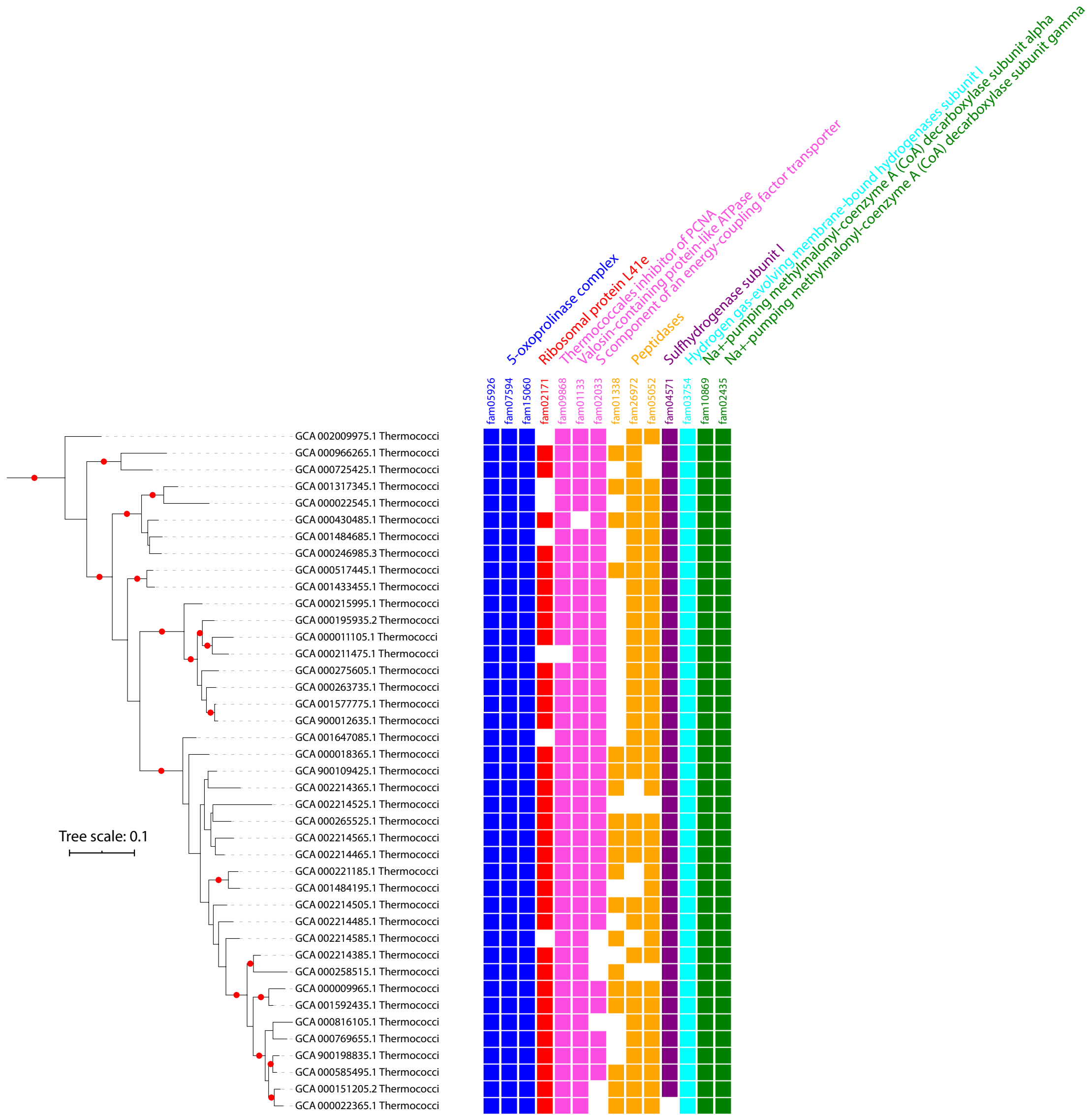


**Supplementary Figure S11.** Presence and absence of 14 families of module 8 in genomes of Thermococci. Scale bar indicates the average substitutions per site.

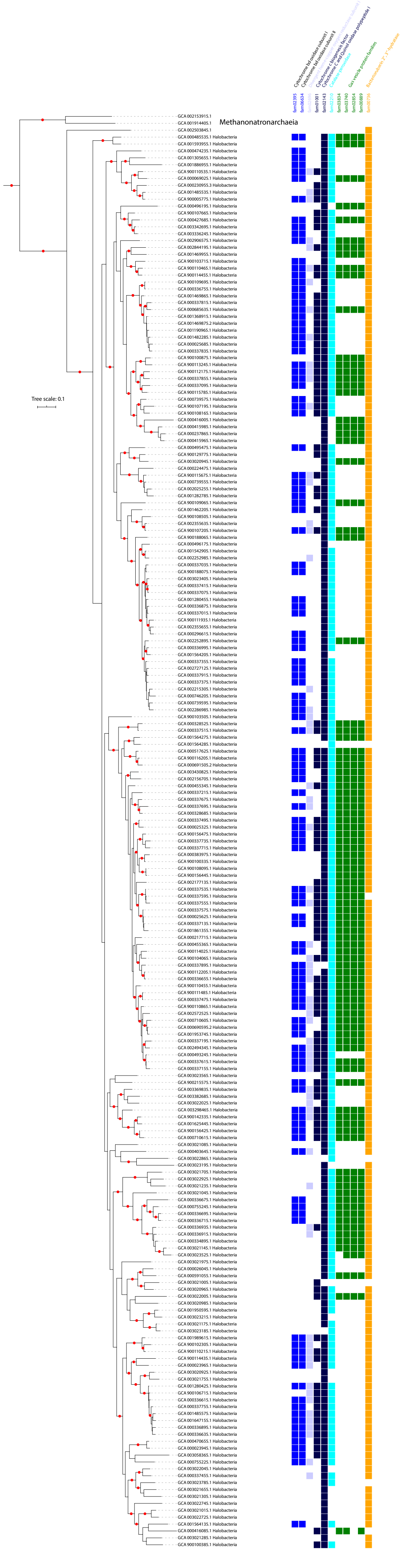
Cytochrome bd oxidase subunit I
Cytochrome bd oxidase subunit II
Divergent heme-copper oxygen reductase subunit I
Cytochrome c biogenesis factor
Cytochrome C and Quinol oxidase polypeptide I
Catalase-peroxidase
Gas vesicle protein families
Bacterioruberin 2'', 3''-hydratase
fam02395
fam06634
fam02696
fam01001
fam02143
fam02210
fam03834
fam03740
fam02854
fam00889
fam00736
Methanonatronarchaeia
Tree scale: 0.1

**Supplementary Figure S12.** Presence and absence of 11 families of modules 13 and 108 in genomes of Halobacteria. Scale bar indicates the average substitutions per site.

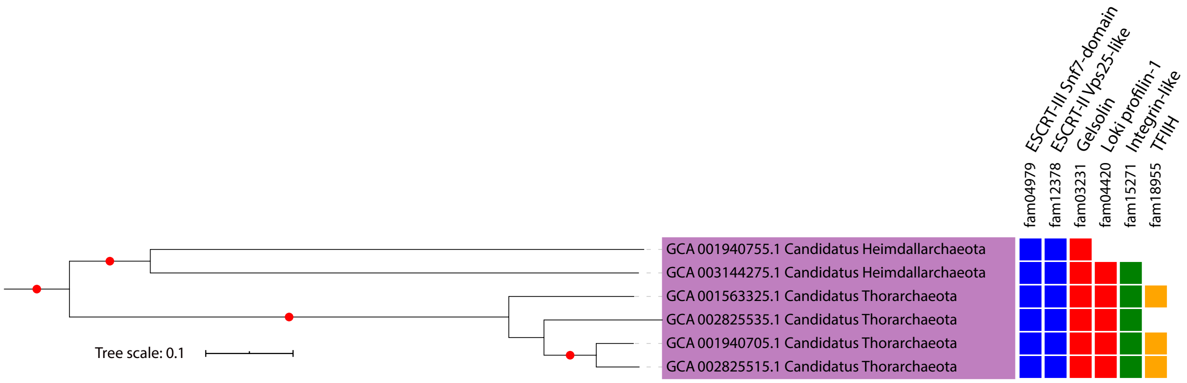


**Supplementary Figure S13.** Presence and absence of 6 families of module 48 in genomes of Asgard. Scale bar indicates the average substitutions per site.

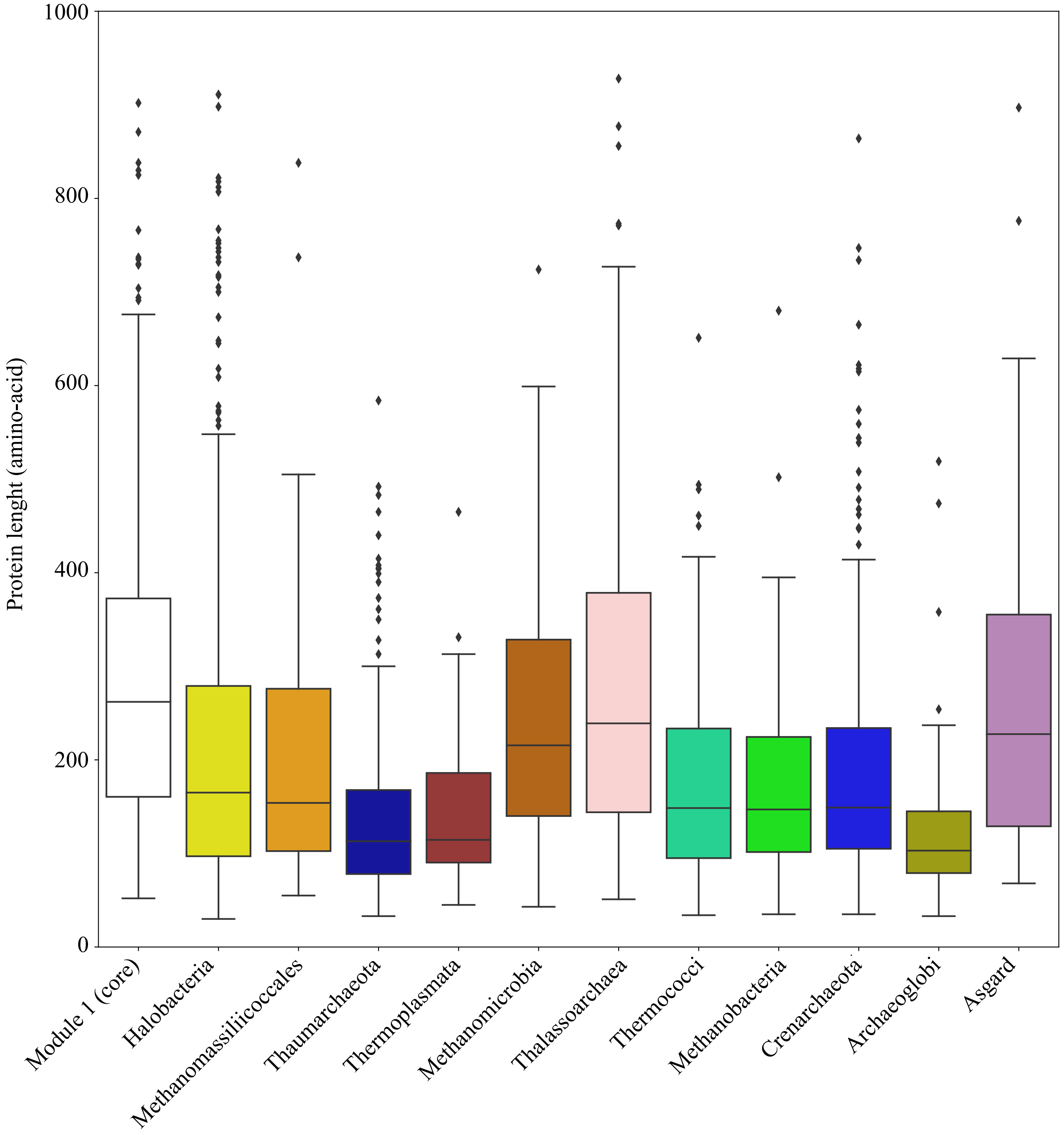


**Supplementary Figure S14.** The length distribution of hypothetical proteins (in amino acid).

**Supplementary dataset**

**Table S1.** List of the 3,197 genomes used in this study. For each genome (column A), the old name, the number of scaffolds, genome size and number of CDS are displayed in columns B, C, D and E. Genome source is in column F, dRep cluster in column G. Genome completeness and the contamination based on single copy genes are displayed in columns H and I respectively. Column J informs about the concatenated ribosomal proteins. The 1,179 representative genomes are indicated in column K. The phylum and superphylum (DPANN and non-DPANN) taxonomy of the representative genomes are provided in columns L and M. Taxonomy based on the different databases we pulled out the genomes is shown in column N.

**Table S2.** Taxonomy distribution of the 113 modules. Module name is indicated in column A whereas the number of families is indicated in column B. Suggested taxonomic distribution is indicated in column C. Column D details the genomes used to define the taxonomic distribution (phylum, number of genomes).

**Table S3.** Annotation of the 10,866 families. Column A: module number. Column B: family accession. Column C: number of proteins in the family. Column D: median length of the proteins. Column E: ratio of proteins predicted to contain a signal peptide. Column F: median number of predicted transmembrane helix per protein. Column G: domain architecture reported by Pfam. Columns H, I, J, K, L: KEGG annotations. Column M: Cazy annotation. Columns N to AC indicate the ratio of genomes having the given family in the given archaeal phylum. Columns AD to CK indicate the ratio of genomes having the given family in the given bacterial phylum.

**Table S4.** Annotation of the subfamilies (column C) based on Hmmsearch against the PDB database (columns D and E) and based on HMM-HMM prediction against the arCogs of the EggNOGs database (columns F, G, H and I).

**Table S5.** Genes neighboring the four genes encoding the subunits of the ammonia monooxygenase. The four genes downstream and upstream of each *amoA*, *amoB*, *amoC* and *amoX* genes (column H) were identified and annotated using the protein clustering (column E), the PFAM (column G) and the KEGG databases (column F).

**Table S6.** Genes neighboring the three genes encoding the subunits of the 5-oxoprolinase complex. The three genes downstream and upstream of each *pxpA*, *pxpB* and *pxpC* genes (column H) were identified and annotated using the protein clustering (column E), the PFAM (column G) and the KEGG databases (column F).

**Table S7.** Genes neighboring the three genes encoding the enzymes of the pathway of the C50 carotenoid bacterioruberin and the gene encoding a distant homolog of the catalytic subunit I of heme-copper oxygen reductase (fam02696). The four genes downstream and upstream of each *LyeJ*, *CruF* ,*CrtD* and fam02696 genes (column H) were identified and annotated using the protein clustering (column E), the PFAM (column G) and the KEGG databases (column F).

**Table S8.** Genes neighboring the two genes encoding the integrin beta 4 and the TFIIH. The five genes downstream and upstream of each *integrin* and the *TFIIH* genes (column H) were identified and annotated using the protein clustering (column E), the PFAM (column G) and the KEGG databases (column F).
